# MYB68 orchestrates cork differentiation by regulating stem cell proliferation and suberin deposition

**DOI:** 10.1101/2024.03.06.583666

**Authors:** David Molina, Sara Horvath, Xudong Zhang, Wei Xiao, Noah Ragab, Dagmar Ripper, Joachim Kilian, Tonni Grube Andersen, Laura Ragni

**Affiliations:** ZMBP-Center for Plant Molecular Biology, University of Tübingen, Auf der Morgenstelle 32, D-72076 Tübingen, Germany; University of Freiburg, Institute of Biology II, Schänzlerstr 1, 79104 Freiburg, Germany; Max Planck Institute for Plant Breeding Research, Carl-Von-Linné-weg 10, 50829, Cologne, Germany

## Abstract

Plants have developed specialized barriers to protect and isolate the inner tissues from the environment while maintaining homeostasis. Different barriers are present in various organs and at different growth stages. During secondary growth, the periderm acts as the protective tissue, covering roots, stems, and branches as they become thick. The periderm is a dynamic barrier comprising a stem cell niche known as the cork cambium, which bifacially divides to generate the phelloderm inward and the cork outward. Cork cells have a unique cell wall impregnated with suberin and lignin polymers, essential for the barrier function.

Despite its importance, the differentiation process that forms new cork cells from the stem cell is largely unknown. In this work, we identify members of the MYB36-subclade transcription factors as key regulators of cork differentiation. On the one hand, this set of transcription factors promotes suberin deposition by inducing the expression of enzymes involved in all steps of suberin biosynthesis, including the recently discovered suberin-polymerizing enzymes GDS Lipases; on the other hand, it represses cork cambium proliferation. Furthermore, we demonstrated that suberin deposition in the cork is a robust process regulated by a complex network of transcription factors, including other MYB transcription factors that activate suberin deposition in the endodermis. However, only members of the MYB36 subclade can repress cell proliferation in different developmental contexts, highlighting general and specific functions for MYB transcription factors. These findings have broad applicability, as tissue-specific manipulation of MYB activity has the potential for improving traits of biotechnological interest, such as thicker periderms and more suberized cork layers, and for assessing how these traits affect plant performance in response to stresses.

## Introduction

Plant barriers protect and isolate the vasculature from the environment, regulating nutrient assimilation, gas exchange, water loss, and pathogen penetration (Serra *et al*., 2022). Depending on the plant family/species and developmental stage, different barriers develop in roots. During primary growth, the endodermis, present in all vascular plants, and the exodermis, occurring in many flowering plants, are the central protective tissues of roots (Barberon *et al*., 2016; Doblas *et al*., 2017; Enstone *et al*., 2002; Serra *et al*., 2022). The periderm replaces the endodermis in the older part of the roots, undergoing extensive radial thickening. This is typical of trees but also occurs in many herbaceous plants, such as Arabidopsis (Serra *et al*., 2022; Wunderling *et al*., 2018). In contrast to the other barriers, the periderm comprises a stem cell niche, the cork cambium, that allows the barrier to follow the radial growth of the organs. The cork cambium can remain active for many years or throughout plant life, producing the cork tissue towards the environment and the phelloderm towards the vasculature (Fig. 1A) (Campilho *et al*., 2020; Evert, 2006; Serra *et al*., 2022).

**Figure 1:**
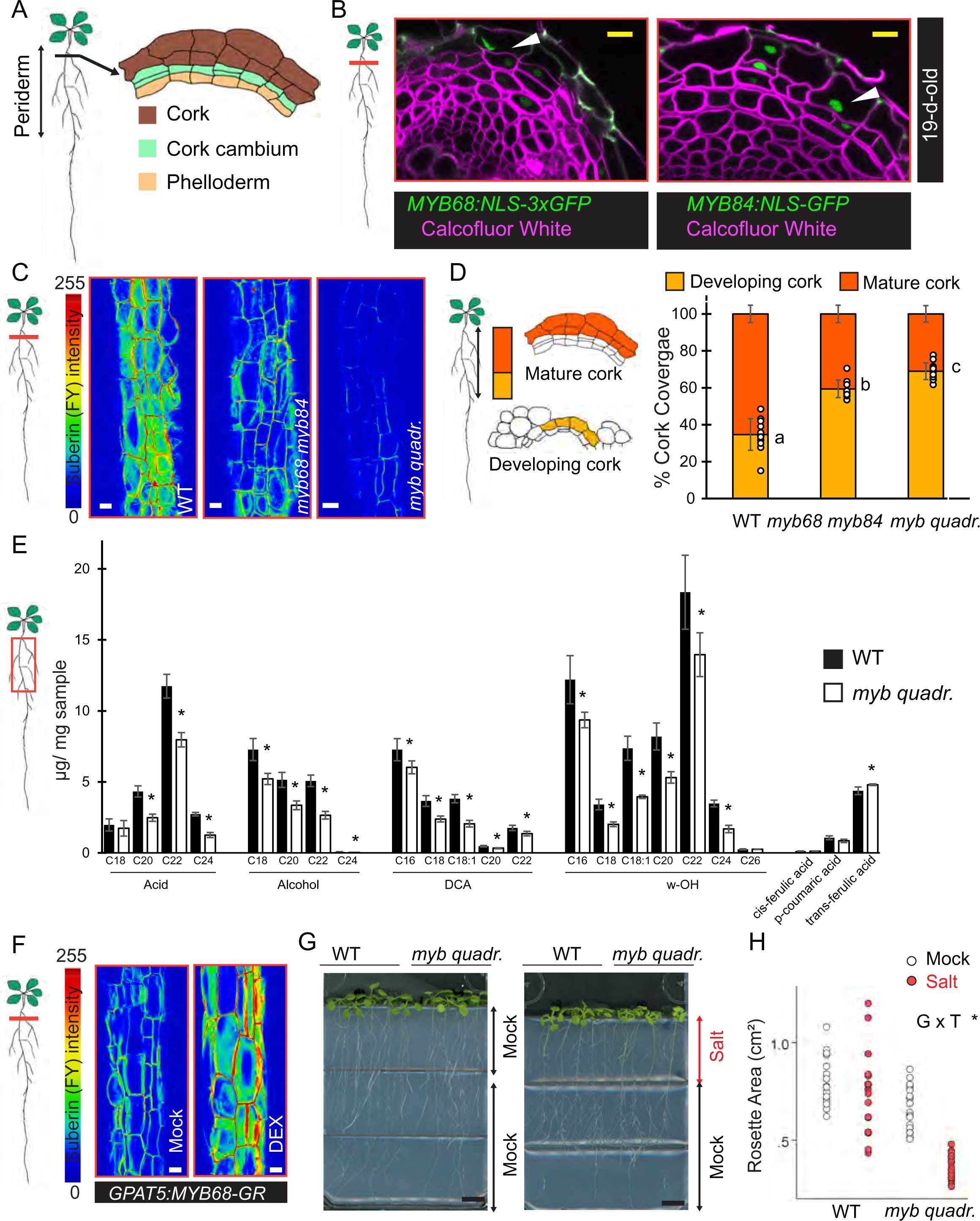
(A) Sketch of a periderm, the different tissue layers are highlighted in distinct colors: cork: brown, green: cork cambium, beige: phelloderm. (B) Confocal microscopy images of vibratome cross-sections of 19-day-old roots showing the expression of *MYB68* (*pMYB68: NLS-3xGFP*; Green) and *MYB84 (pMYB84:NLS-3xGFP*; Green) in the cork (white arrows). The cells are outlined by Calcofluor White staining (magenta). (C and D) Suberin quantification by Fluorol yellow (FY) staining and suberin coverage in the cork of wild type (WT), *myb68 myb84*, and *myb quadr.* (*myb37 myb38 myb84 myb68*) roots. (C) Representative images showing relative intensities of FY fluorescent signal (14-day-old plants). (D) Suberin deposition in the Arabidopsis root periderm can be divided into two distinct stages. In the Developing cork (pale orange) region, only a few cork cells are suberized, while in the mature cork (dark orange) region, all cork cells are already differentiated. Relative quantification of developing and mature cork suberin regions in 19-day-old plants. (E) Quantification of suberin constituents (µg/ mg dry sample) via GC-MS in the cork of WT and *myb quadr*. Suberin constituents include DCA (α,ω-dicarboxylic acids), ω-OH (ω-hydroxy fatty acids), Alcohol (fatty alcohols), acid (fatty acids), and aromatic compounds (cis-ferulic Acid, p-coumaric Acid and trans-ferulic Acid). (F) Representative images showing fluorescence relative intensities of FY signal in the cork of pGPAT5:MYB68-GR, induced and not induced roots. 9-day-old plants were treated for 5 days with mock or 10µM DEX. (G and H) Salt effect on *myb quadr.* and WT growth. (G) Plants were grown for 9 days prior to transfer to split plates (Most upper part is supplemented with 150 mM NaCl or mock) for 5 days. H) Quantification of the rosette area (cm²) of (G). In (D) One-way ANOVA (95% CI, post hoc: Tukey HSD, n = 10–14). In (E) Student T-Test (*: P<0.05, n:3-4). In (H) Two-way ANOVA, G*T: Genotype (WT or *myb quadr*.) Treatment (mock or WT) (95% CI, *:P<0.05, n = 16-18). Black bars: 1cm, white scale bars: 20µm, yellow scale bars: 10µm.

Common to all root barriers is the presence of specialized polymers such as suberin and lignin that impregnate the cell walls (Serra *et al*., 2022). The endodermis is characterized by localized lignin depositions known as "Casparian strips" and suberin lamellae, which accumulate across the entire plasma membrane (Doblas *et al*., 2017; Naseer *et al*., 2012). Similarly, the exodermis is a ligno-suberized layer in which lignin deposition occurs in a U-shaped fashion before suberin accumulation (Cantó-Pastor *et al*., 2024; Manzano *et al*., 2022). The cork cell wall is also impregnated with both polymers (Kosma *et al*., 2015; Serra *et al*., 2022; Wunderling *et al*., 2018). Plants with lower suberin contents in the endodermis and or periderm are less resistant to abiotic stresses, highlighting the protective function of these polymers, while periderm thickness correlates with the ability to withstand wildfire and pathogen penetration (Andersen *et al*., 2021; Barberon *et al*., 2016; Graves *et al*., 2014; Shukla *et al*., 2021).

The recalcitrant nature of suberin, which is even more resistant than lignin in soil (Harman-Ware *et al*., 2021), has sparked great interest in the molecular regulation of polymer accumulation in root barriers for biotechnological purposes (Harman-Ware *et al*., 2021). The molecular network underlying the different steps of endodermis differentiation is well characterized, and different transcription factors regulate the specific steps. MYB36 serves as the master regulator of Casparian strip formation, directly activating *PEROXIDASES* (*PERs*), *RESPIRATORY BURST OXIDASE HOMOLOGs* (*RBOHs*) and scaffold proteins such as Casparian strip membrane domain proteins (CASPs), and *DIRIGENT PROTEINs* (DIRs) required for polar lignin deposition (Gao *et al*., 2023; Kamiya *et al*., 2015; Liberman *et al*., 2015; Wang *et al*., 2022). Suberin deposition in the endodermis is promoted by a large array of MYB TFs belonging to different subclades (Dubos *et al*., 2010), such as the S10 (MYB9, MYB39, MYB107), the S11 (MYB41, MYB74) and S24 subclade (MYB53, MYB92, MYB93), (Cohen *et al*., 2020; Kosma *et al*., 2014; Lashbrooke *et al*., 2016; Shukla *et al*., 2021; Wang *et al*., 2020; Xu *et al*., 2022). Suberin is highly dynamic, and several hormones, such as abscisic acid (ABA), ethylene, and gibberellins, determine suberin homeostasis (Barberon *et al*., 2016; Binenbaum *et al*., 2023; Wei *et al*., 2019). For instance, ABA is well-known to trigger suberin deposition via the activation of MYBs (Shukla *et al*., 2021; Wei *et al*., 2019), while ethylene application and iron deficiency have a repressive effect on suberin deposition (Barberon *et al*., 2016).

Less is known about periderm ontogenesis and cork differentiation. Recently, we demonstrated that auxin is required for the initial formative divisions in the pericycle, which gives rise to the cork cambium; in addition, auxin is necessary to maintain an active cork cambium (Xiao *et al*., 2020). Downstream of auxin, the transcription factors WUSCHEL RELATED HOMEOBOX 4 (WOX4) and BREVIPEDICELLUS (BP)/KNAT1, promote cork cambium proliferation (Xiao *et al*., 2020). However, the differentiation process of the cork cambium cells into cork cells and the regulation of suberin deposition in the cork remain unknown. QsMYB1 (ortholog of MYB84/MYB68 from cork oak (*Quercus suber*)) has been proposed to control cork differentiation (Almeida *et al*., 2013; Capote *et al*., 2018). However, functional characterization is still missing due to limitations in genetic analysis with cork oak trees. Interestingly, MYB84 has been previously employed as a periderm marker in Arabidopsis root, paving the way for functional studies (Wunderling *et al*., 2018; Xiao *et al*., 2020). MYB84 and MYB68 belong to the S14 subclade of R2-R3 MYB transcription factors (from now on referred to as the MYB36 subclade)(Dubos *et al*., 2010), which includes MYB36. Here, we explored the role of the MYB36 clade transcription factors during cork differentiation. We demonstrated that MYB68 and, to a lesser extent, other members of the clade regulate suberin deposition in the cork via the activation of suberin polymerizing and biosynthesis genes. Moreover, induction of MYB68 in the cork is sufficient to increase suberin levels, while induction in the periderm results in plants with an extra cork layer, traits of biotechnological interest in programs aiming to improve soil carbon sequestration. Interestingly, this set of MYBs also represses cork cambium proliferation. This stem cell repressive differentiation behavior is limited to the MYB36 subclade, while MYBs from neighboring clades also contribute to regulating cork suberin levels, highlighting that suberin deposition in the cork is a robust process controlled by multiple sets of MYB transcription factors.

## Results

### All members of the MYB36 subclade are expressed during periderm development

The MYB36-containing subclade (S14 subclade) comprises six members: MYB36, MYB37, MYB38, MYB68, MYB84, and MYB87. We first focused on MYB68 and MYB84, which represent the closest orthologs of QsMYB1 in Arabidopsis, and mapped their expression pattern during the early and late steps of periderm development by employing promoter-fluorophore reporter lines. Both MYB promoters were active during periderm formation (Fig. S1A), and their expression was maintained in the periderm of old roots (19-day-old) (Fig. 1B and S1B). MYB68 was expressed predominantly in the cork cambium and a few cork cells, whereas the expression of MYB84 was broader toward the cork (Fig. 1B and S1A). Consistently, MYB68 and MYB84 signals overlapped with the suberin biosynthesis gene reporter *GLYCEROL-3-PHOSPHATE-SN-2-ACYLTRANSFERASE 5* (*GPAT5*) (Wunderling *et al*., 2018) in the cork, with a similar *PER15* reporter (Xiao *et al*., 2020) in early differentiating cork and cork cambium and with WOX4 (Xiao *et al*., 2020) in the cork cambium (Fig. S1C). Notably, this expression pattern is consistent with a role for MYB68 and MYB84 in the early steps of cork differentiation. Interestingly, we found that the other members of the S14 subclade were also expressed during periderm formation in overlapping and distinct domains. *MYB37/REGULATORY AXILLARY MERSITEM 1 (RAX1)* and *MYB38/RAX2* expression encompassed all periderm layers, while *MYB36* was restricted to the mature cork, and *MYB87* to the cork cambium. Overall, our findings suggest that the MYB S14 subclade contributes to periderm formation and differentiation.

### MYB68 and other MYB36 subclade TFs promote suberin deposition in the cork

The hallmark of cork differentiation is suberin deposition (Serra *et al*., 2022), which can be easily imaged by fluorescent staining. Fluorol yellow (FY) staining, compared to analytical approaches, is a quick method to quantify suberin deposition and allows spatial resolution (Andersen *et al*., 2021; Lux *et al*., 2005; Ursache *et al*., 2018; Wunderling *et al*., 2018). Thus, to investigate the role of S14-subclade MYBs in suberin deposition, we performed FY staining of loss-of-function mutants. In the mature cork of roots of *myb68, myb84*, *myb87*, *myb36* single mutants, *myb68 myb87* double mutant, and *myb37 myb38 myb84* triple mutants, we observed normal suberin deposition (Fig. S1D-E, S2A) suggesting redundancy among the different members of clade S14. In agreement, the inactivation of both *MYB68* and *MYB84* led to a mild reduction of suberin, while only in the *myb37 myb38 myb68 myb84* quadruple mutant (*myb quadr.*) we detected a drastic decrease in suberin deposition in the cork (Fig. 1C).

Interestingly, when we explored the suberin deposition pattern in the whole cork, we noticed a reduction of the mature cork region compared to the developing-cork region in *myb68* single mutants, while *myb87*, *myb84*, *myb37 myb38 myb84,* and *myb36* did not show any phenotypes (Fig. S2B-D). *myb68 myb87* double mutants were indistinguishable from *myb68* single mutants, whereas *myb68 myb84* double mutants and the *myb* quadruple mutants showed a more substantial reduction than *myb68* single mutants (Fig. 1D, S2C). Importantly, these observations cannot be explained by impaired primary root growth or delay in periderm development in *myb* mutants as they display root lengths comparable to WT (or even slightly longer) and slightly increased cork-to-root -length ratio, suggesting normal primary growth and periderm development (Fig. S2E-F). Altogether, the analysis of MYB loss-of-function mutants suggests that MYB68 and, to a lesser extent, MYB84, MYB37, and MYB38 contribute to cork differentiation. Consistent with a key role for MYB68 in promoting suberin deposition, cork-specific induction of MYB68 (*pGPAT5:MYB68-GR*) was sufficient to rescue the *myb68 myb84* suberin phenotypes (Fig. S2G-H).

Suberin is a complex polymer based on ester-bond linked glycerol, fatty acids, their oxidized derivates, and ferulic acid (Serra and Geldner, 2022). Recent work showed that although ferulic acid represents only a minor fraction of the total suberin-related components, it is crucial for de novo suberin deposition (Andersen *et al*., 2021). We performed Gas Chromatography–Mass Spectrometry (GC-MS) analysis to investigate whether the chemical composition of suberin is affected in the *myb37 myb38 myb68 myb84* quadruple mutant. In the *myb37 myb38 myb68 myb84* quadruple mutant, we observed a reduction of most of the aliphatic components compared to WT, while aromatic acids, such as ferulic acid, were unchanged or slightly increased, indicating that MYB68, MYB84, MYB37, and MYB38 regulate the fatty acid suberin biosynthesis pathway (Fig. 1E) rather than the phenylpropanoid pathway, source of ferulic acid. Monolignols, lignin building blocks, are also produced by the phenylpropanoid pathway. In agreement with this, we did not observe any decrease of lignin accumulation in the cork of the *myb68 myb84* double mutants and *myb* quadruple mutants (Fig. S2I). Next, we wondered if MYB68 is sufficient to promote suberin deposition in the cork. To tackle this, we generated tissue-specific inducible lines using the *GPAT5* promoter, which is strongly expressed in the cork during periderm development (Leal *et al*., 2022; Wunderling *et al*., 2018). Upon induction of MYB68 in the cork, we detected an increase of FY signal in the cork, proving that MYB68 is sufficient to trigger suberin accumulation (Fig. 1F).

Reduced suberin deposition impacts barrier functionality and permeability, and thus, suberin mutants are less resistant to salt stress (Andersen *et al*., 2021; Ursache *et al*., 2021). We assessed the physiological impact of reduced suberin deposition of *myb quadr*. mutants, by exposing the root region encompassing the periderm to salt stress (by employing split plates). This is a fundamental requirement to study periderm-specific responses as salt stress strongly impacts primary root growth and suberin accumulation in the endodermis (Andersen *et al*., 2021; Barberon *et al*., 2016; Duan *et al*., 2015; Julkowska *et al*., 2014; Ursache *et al*., 2021), which may lead to indirect effects in the periderm. Primary root length in treated or untreated plants was comparable (Fig. 1G); however, the shoots of the *myb37 myb38 myb68 myb84* quadruple mutants were smaller and more compact in the salt-treated plants, showing that impaired cork differentiation affects tolerance to salt (Fig. 1G-H).

### *MYB68* promotes the expression of suberin biosynthesis and polymerizing enzymes

To explore how these MYBs regulate suberin deposition, we profiled the transcriptome of *myb68 my84* double mutants in comparison to their parental backgrounds. To avoid developmental heterogeneity, we sampled only the uppermost two cm of 18-day-old roots, representing a uniform growth stage with a fully established periderm. Among the downregulated genes (Log_2_FC <-0.5) in *myb68 myb84*, we found genes coding for well-established suberin biosynthesis enzymes, such as *CYTOCHROME P450 FAMILY 86* (*CYP86A1)/HORST*, *CYP86B1/RALPH*, *GPAT5*, *FATTY ACID REDUCTASEs* (*FARs*), *3-KETOACYL-COENZYME A SYNTHASE* (*KCSs*)(Fig. S3A), which are expressed in the cork (Leal *et al*., 2022; Serra *et al*., 2022; Wunderling *et al*., 2018). In this group, we also noticed genes encoding for the recently characterized endodermal GDSL-type esterase/lipase proteins (GELP38, GELP51, GELP49, and GELP96), which polymerize suberin in the apoplast (Ursache *et al*., 2021). In accordance with the idea that GELPs may also act in the cork, many GELPs that are downregulated in *myb68 myb84* are enriched in cork tissue *versus* whole-root tissues (Leal *et al*., 2022). By employing fluorescent reporters, we confirmed that *GELP38*, *GELP51*, *GELP96,* and *GELP49* are expressed in the cork (Fig. 2A). Furthermore, the loss-of-function of five GELPs (*gelp22 gelp38 gelp51 gelp49 gelp96: gelp q.*) had no detectable FY signal in the periderm (Fig. 2B-C), pointing out that GELPs are also required for suberin polymerization in the cork. Notably, the expression of GELP51 in the cork (*pGPAT5:GELP51*) of *myb68 myb84* resulted in the restoration of the suberin levels similar to WT (Fig. 2D). Additionally, the defects in the suberin deposition pattern of *myb68 myb84* were partially rescued by the expression of GELP51 (Fig. 2E), highlighting the importance of GELP enzymes in suberin deposition.

**Figure 2:**
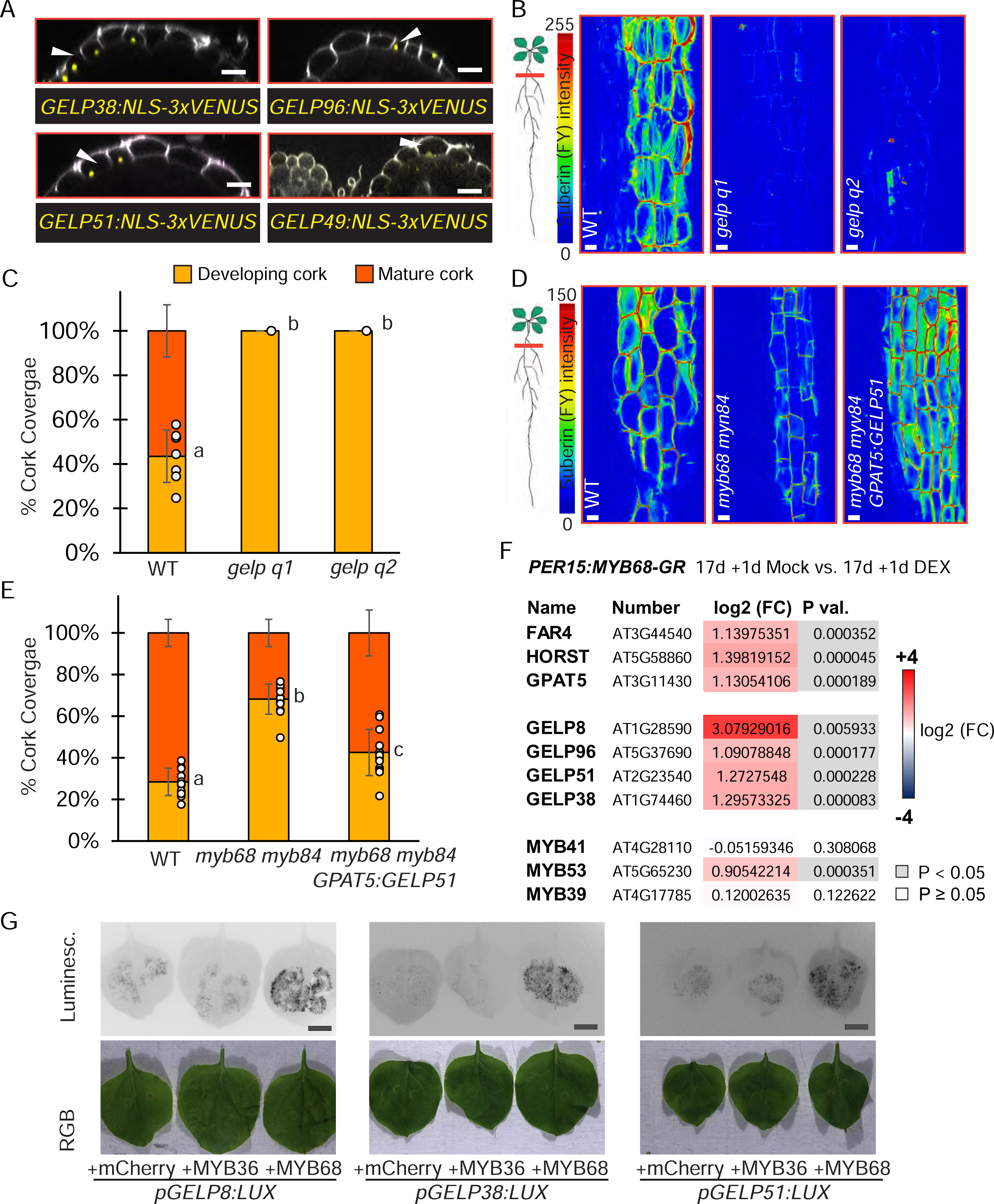
(A) Orthogonal view of z stacks showing the expression of GELP38 (*pGELP38:NLS-3xVenus*), GELP96 (*pGELP96:NLS-3xVenus*), GELP51(*pGELP51:NLS-3xVenus*) and GELP49 (*pGELP49:NLS-3xVenus*) in the cork (white arrows) (14 day-old plants). (B) Representative images showing relative intensities of FY fluorescent signal in the cork of WT, *gelp q1* and *gelp q2* (*gelp38 gelp51 gelp49 gelp96 gelp22*) (14 day-old-plants). (C) Relative quantification of developing and mature cork suberin regions of WT, *gelp q1,* and *gelp q2* 19-day-old roots. (D) Representative images showing relative intensities of FY fluorescent signal of 14-day-old WT, *myb68 myb84,* and *pGPAT5:GELP51 myb68 myb84* roots. (E) Relative quantification of developing and mature cork suberin regions of 19-day-old WT, *myb68 myb84,* and *pGPAT5:GELP51 myb68 myb84* plants. (F) Log2 Fold Change (FC) of relative expression of selected suberin biosynthesis, polymerizing, and regulating genes upon induction of MYB68 in the periderm (*pPER15:MYB68-GR;* plants were grown 17 days prior to treatment with mock or 10 µM DEX for 24h.). Data were obtained by qPCR, as described in the methods section. p values (P val.) are based on the Student’s t-test mock vs DEX. (G) Transactivation studies in *N. benthamiana* leaves using luciferase reporting system. Upper panels show the luminescence intensities of leaves co-infiltrated with an effector construct (*p35S:mCherry*, *p35S:MYB68-mCherry*, *p35S:MYB84-mCherry*), a reporter construct (promoter of interest fused with luciferase (LUX) reporter (*pGELP8:LUX*, *pGELP38:LUX* and *pGELP51:LUX*), and the FBP11 construct (encoding the fungal bioluminescent pathway. Lower panels show the RGB images of the same leaves. In (C,E) One-way ANOVA (95% confidence interval [CI], post hoc: Tukey HSD, n = 10-14). Black scale bars: 1cm and white scale bars: 20µm.

Interestingly, the expression of other MYB TFs belonging to the MYB41- and MYB39-containing subgroups (subclade S11 and S10, respectively), which are known to control suberin deposition in the root endodermis and in the seed coat (Cohen *et al*., 2020; Kosma *et al*., 2014; Lashbrooke *et al*., 2016; Shukla *et al*., 2021), was unchanged or increased in the *myb68 myb84* dKO with exception for MYB53. This suggests that MYB68 directly impacts aliphatic suberin biosynthesis and polymerization.

To confirm that MYB68 activates the expression of suberin-associated genes in the cork, we exploited a complementary approach: we shortly induced MYB68 in the periderm (*pPER15:MYB68-GR*) and checked the expression of a set of genes by qPCR. *FAR4, HORST/CYP86A1, GPAT5, GELP8, GELP51, GELP96, GELP38 and MYB53* were significantly upregulated upon MYB68 induction indicating that MYB68 triggers the suberin pathway at multiple steps (Fig. 2F). In agreement, the expression of MYB68 in tobacco leaves triggered the activation of luminescent reporters for suberin biosynthesis enzymes (*pGPAT5:nnLUX*, *pFAR4:nnLUX*; Fig. S3B-C), suberin polymerizing enzymes (p*GELP8:nnLUX*, *pGELP38:nnLUX*, *pGELP51:nnLUX*; Fig. 2G) and suberin regulatory genes (*pMYB53:nnLUX*; Fig S3D) suggesting that MYB68 directly activate them. Interestingly, MYB36 was able to promote only the expression of *pFAR4:nnLUX* and *pCASP2:nnLUX* (CASP2 is a known MYB36 target (Kamiya *et al*., 2015)) (Fig. S3E), while MYB68 could not activate the expression of *pCASP2:nnLux* (Fig. S3B,E) highlighting the specificity among these TFs.

Altogether, these results show that MYB68, but not MYB36, has the ability to activate suberin deposition in the cork.

### A specific set of MYBs represses stem cell proliferation in the cork cambium

The observation that the expression pattern of MYB68, MYB84, MYB37, MYB38, and MYB87 encompasses the unsuberized cork cambium (Fig. S1A-C), prompted us to investigate whether these TFs play additional roles during periderm development. Thus, we investigated whether cork cambium activity is affected in loss-of-function mutants. As a proxy for the cork cambium proliferation rate, we quantified the number of periderm cells per cross-section, while for the cork cambium differentiation rate, the number of cork cells per number of peridermal cells (cork ratio). *Myb84*, *myb87,* and *myb37 myb38 myb84* triple mutants did not display any difference in the total number of periderm cells and cork ratio when compared to WT (Fig. S4A-F). *Myb68* single mutants, *myb68 myb87,* and *myb68 myb84* double mutants showed impaired cork differentiation (decreased cork ratio compared to WT). In *myb68 myb84*, we also observed a trend toward more periderm cells (significant in 6 independent experiment repetitions out 8) while *myb37 myb38 myb68 myb84* quadruple mutants showed stronger phenotypes in comparison to *myb68 myb84* double mutants (FIG. 3A-C). Altogether, these genetic analyses indicate that MYB68 and, to a lesser extent, MYB84, MYB37, and MYB38 repress stem cell proliferation in the cork cambium. Interestingly, in *myb68 myb84* dKO and *myb37 myb38 myb68 myb84 qKO* mutants, we observed gaps in the autofluorescence signal coming from the aromatic components present in differentiated cork cells compared to WT (Fig. S4G; red arrowheads), further highlighting the role of these MYBs in promoting cork differentiation and repressing stem cell proliferation.

**Figure 3:**
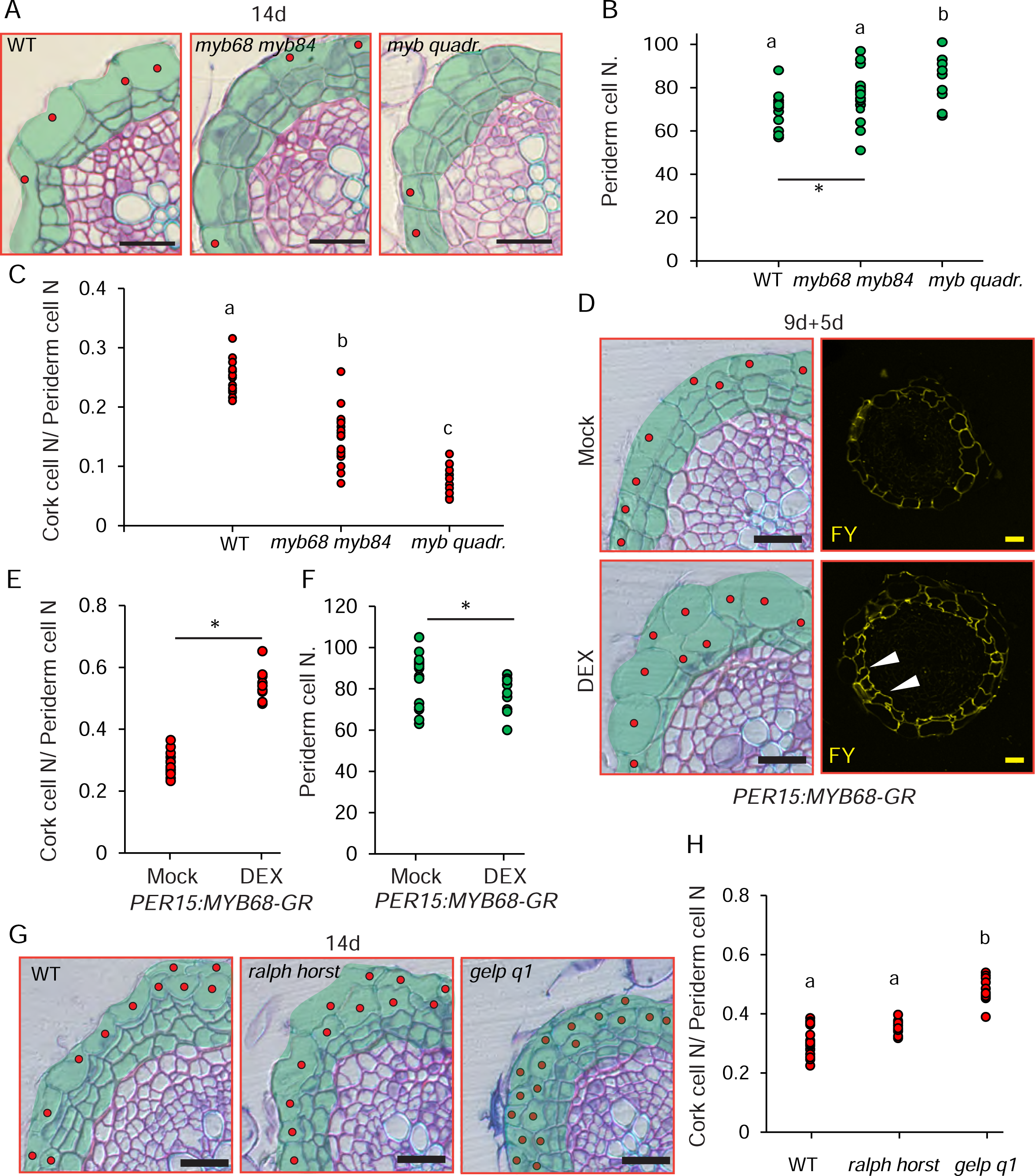
(A) Cross-sections (plastic embedding) of the uppermost part of 14-day-old WT, *myb68 myb84*, and *myb quadr.* roots. (B) Quantification of the total number of periderm cells in the experiment presented in (A). (C) Quantification of the ratio of the number of cork cells/number of periderm cells in the experiment presented in (A). (D) Cross-sections and Fluorol Yellow staining of the uppermost part of 14-day-old *pPER15:MYB68-GR* roots. 9-day-old plants were treated for 5 days with mock or 10 µM DEX. The white arrows highlight the two layers of cork. (E-F) Ratio of the number of cork cells/number of periderm cells and the total number of periderm cells in the experiment presented in (D). (G) Cross-sections of the uppermost part of 14-day-old WT, *ralph horst*, and *gelp q1* roots. (H) Quantification of the ratio of the number of cork cells/number of periderm cells in the experiment presented in (G). In (B-C,H) One-way ANOVA (95% confidence interval [CI], post hoc: Tukey HSD, n = 15-20). In (E-F) Student T-Test ( *: P<0.05, n:15-20). Black and yellow scale bars: 20µm. Red dots indicate cork cells, and the whole periderm is highlighted in green.

In agreement, periderm-specific induction of MYB68 (*PER15:MYB68-GR*) resulted in a decrease in the total number of periderm cells, a higher number of cork cells per total periderm cells, and the formation of an extra suberized cork layer, confirming a dual role for MYB68 in promoting differentiation and repressing proliferation of the cork cambium (Fig. 3D-F).

To exclude that the effect of MYB68 on stem cell proliferation is an indirect consequence of altered suberin content in the cork, we analyzed lines with reduced suberin deposition in the cork for cell proliferation phenotypes. The suberin biosynthesis mutant *ralph horst* (Salas-González *et al*., 2021), which mimics *myb68 myb84* mutants regarding FY staining (Fig. S4H-I ), was undistinguishable from WT in terms of periderm cell number and cork per total number of periderm cells (Fig. 3G-H). In addition, *gelp* quintuple mutants, which almost lack FY the cork, displayed an increase of cork per periderm cell ratio and reduced periderm cell number (Fig. 3G-H), which is the opposite phenotype of *myb37 myb38 myb68 myb84* mutants supporting the idea that this group of MYBs plays a specific function in regulating stem cell proliferation.

### MYB68 and MYB84 regulate stem cell proliferation via WOX4 and BP

Auxin plays a pivotal role in orchestrating cork cambium initiation and maintenance (Xiao *et al*., 2020), which begs the question of whether the repressive function of MYB68, MYB84, MYB37, and MYB38 in the cork cambium is dependent on the auxin signaling network of the periderm. To tackle this, we took advantage of the fact that blocking polar auxin transport via pharmacological interference (N-1-naphthylphthalamic acid (NPA)) results in delayed periderm formation/growth (Xiao *et al*., 2020) and tested whether NPA treatment suppresses the periderm overproliferation phenotype of *myb37 myb38 myb68 myb84* quadruple mutants. NPA treatment in WT decreased periderm cell numbers, while in *myb quadruple* mutants, no effect was measurable (Fig. 4A-C). Mechanistically, there could be two alternative explanations: these MYBs act in independent pathways, or crosstalk with auxin-induced periderm program occurs downstream of the auxin signaling perception machinery. To disentangle this, we tested whether MYB68 and MYB84 genetically interact with WOX4 and BP, two known cork cambium regulators downstream of auxin signaling (Xiao *et al*., 2020). For this purpose, we obtained *wox4 myb68 my84* and *bp myb68 myb84* triple mutants.

**Figure 4:**
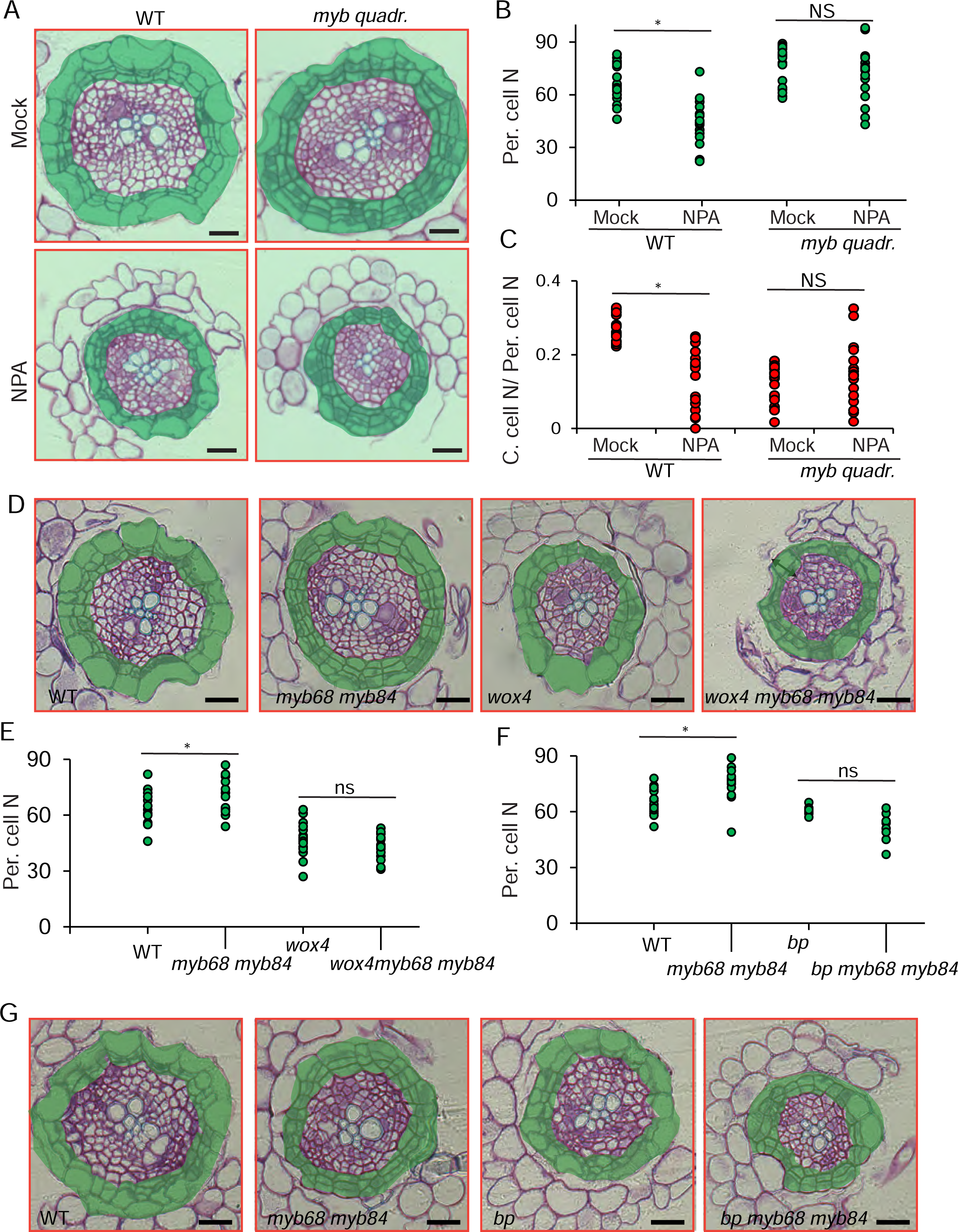
(A) Cross-sections (plastic embedding) of the uppermost part of 14-day-old WT and *myb quadr.* roots treated with N-1-naphthylphthalamic Acid (NPA). 9-day-old plants were treated for 5 days with mock or 10μM NPA. (B-C) Quantification of the ratio of the number of cork cells/number of periderm cells and the total number of periderm cells in the experiment presented in (A). (D) Cross-sections of the uppermost part of 14-day-old WT, *myb68 myb84*, *wox4* and *wox4 myb68 myb84.* (E) Quantification of the total number of periderm cells in the experiment presented in (D). (F) Cross-sections of the uppermost part of 14-day-old WT, *myb68 myb84*, *bp* and *bp myb68 myb84.* (G) Quantification of the total number of periderm cells of the experiment is presented in (F). In (B-C, E-F) Two-tailed Students T-Test ( *: P<0.05, n:15-20). Black scale bars: 20µm. The whole periderm is highlighted in green.

*Wox4* and *bp* mutants showed a reduction in the total periderm cell number compared to WT, confirming previous findings (Xiao *et al*., 2020) (Fig. 4D-G). The inactivation of BP in *myb68 myb84* resulted in a reduction of the total number of periderm cells compared to *myb68 my84* double mutants (Fig. 4F-G). Similarly, the loss of function of *WOX4* in *myb68 my84* led to a decrease in the number of periderm cells (Fig. 4D-E). These results show that WOX4 and BP are necessary for the increased proliferation rate/periderm cell number of the *myb68 myb84*. Interestingly, WOX4 expression was unchanged in *myb68 myb84* double mutant, and BP was only mildly upregulated, suggesting that MYB68 and MYB84 do not directly repress WOX4 and BP expression (Fig. S3A).

### Several sets of MYBs regulate suberin deposition in the cork, but stem cell repression is restricted to the MYB36 clade

We next wondered whether MYBs belonging to subclades that regulate suberin in other developmental contexts (Cohen *et al*., 2020; Kosma *et al*., 2014; Lashbrooke *et al*., 2016; Shukla *et al*., 2021; Wang *et al*., 2020; Xu *et al*., 2022), also play a role in the cork. By exploiting available transcriptomic data for cork-enriched expression (Leal *et al*., 2022) and fluorescent reporters, we found *MYB41*, *MYB53*, *MYB92,* and *MYB93* to be expressed in the cork (Fig. 5A and S3A). Thus, we analyzed suberin deposition in the *myb41 myb53 myb92 myb93* quadruple mutant. The combined loss of function in *MYB41*, *MYB53*, *MYB92*, and *MYB93* resulted in decreased suberin accumulation in the cork and in a reduction of the relative mature-cork region compared to the developing-cork region (Fig. 5B-C), highlighting that suberin deposition in the cork is regulated by several groups of MYBs belonging to different subclades. However, *myb41 myb53 myb92 myb93* quadruple mutants did not show an increased number of periderm cells, reduced cork-to-periderm cell ratio, and gaps of undifferentiated cork as observed in *myb quadr*. mutants, suggesting that they do not repress stem cell proliferation in the cork cambium (Fig. 5D-F).

**Figure 5.**
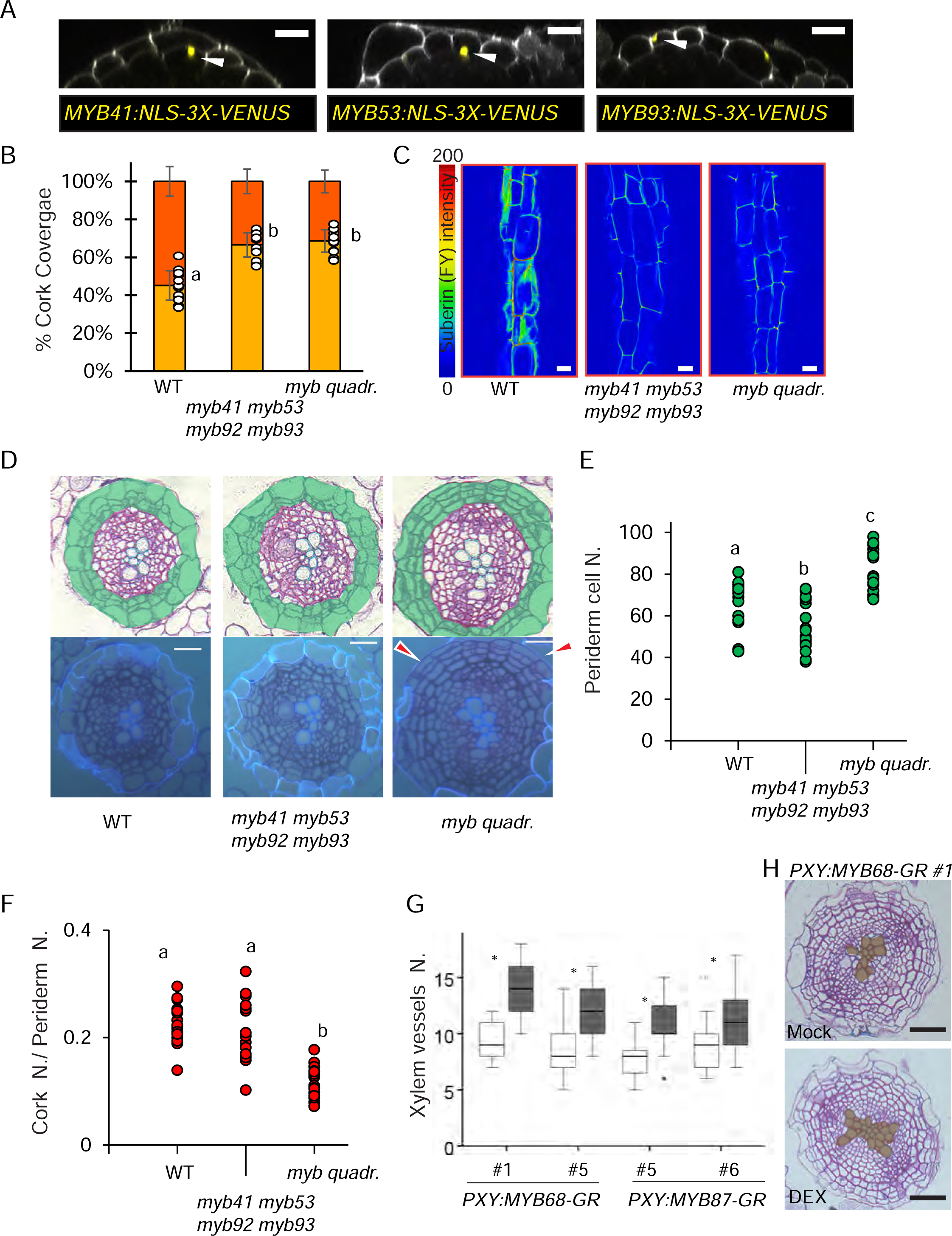
(A) Orthogonal view of z stacks showing the expression of MYB41 (*pMB41:NLS-3xVenus*), MYB53 (*pMYB53:NLS-3xVenus*), and MYB93 (*pMYB93:NLS-3xVenus*) in the cork (white arrowheads) of 14 day-old plants. (B-C) Suberin quantification by Fluorol yellow (FY) staining and suberin coverage in the cork of WT, *myb41myb53 myb92 myb93,* and *myb quadr.* roots. (B) Relative quantification of developing and mature cork suberin regions of 19-day-old plants. (C) Representative images of the relative intensity of FY fluorescent signal (14-day-old plants). (D) Cross-sections (plastic embedding) of the uppermost part of 14-day-old WT and *myb41 myb53 myb92 myb93* and *myb quadr.* roots. The upper panels show bright field pictures, and the lower panels show autofluorescence (DAPI filter) cork cells under UV light. The red arrowheads indicate the absence of differentiated cork cells. (E-F) Quantification of the ratio of the number of cork cells/number of periderm cells and the total number of periderm cells of the experiment presented in (D). (G) Quantification of the number of xylem vessels upon induction of MYB68 and MYB87 in the vascular cambium (*pPXY:MYB68-GR* and *pPXY:MYB87-GR*; 9-day-old plants were treated for 5 days with mock or 10μM DEX) in the cross-sections shown in (H and S4J). (H) Cross-sections of the uppermost part of 14-day-old *pPXY:MYB68-GR* roots. In (B, E-F) One-way ANOVA (95% confidence interval [CI], post hoc: Tukey HSD, (B) n = 10-14, (E-F) n=15-20. In (G) Student T-Test ( *: P<0.05, n:15-20). Black scale bars: 20µm. The whole periderm is highlighted in green.

Finally, we investigated whether MYB68 can promote differentiation and repress cell proliferation in other developmental contexts. As MYB87 has been previously described as a repressor of vascular cambium activity and overexpression of MYB87 leads to reduced radial growth (Zhang *et al*., 2019), we tested whether MYB68 can repress vascular cambium proliferation.

Vascular cambium-specific overexpression of both MYB87 and MYB68 (*PXY:MYB68-GR* and *PXY:MYB87-GR*) resulted in an increase of xylem vessel numbers due to enhanced differentiation and reduced proliferation of the vascular cambium, showing that MYB68 can repress vascular cambium activity (Fig. 5G-H, S4J). Altogether, these findings indicate that multiple MYB transcription factors belonging to different subclades regulate suberin deposition in the cork, while only MYBs of MYB36-sub clade repress stem cell proliferation.

## Discussion

The plant body is protected from biotic and abiotic stress by different barrier tissues depending on the organ, plant species, and age. The periderm, the protective tissue of the organs undergoing radial thickening, has been extensively described in many plant species. However, it has only recently started to be characterized at the molecular level thanks to advances in omics technologies and the establishment of the Arabidopsis as a model for studying periderm development and cell biology (Wunderling *et al*., 2018). The periderm is a dynamic barrier, including a stem cell population, which can remain active for many years or throughout the entire plant life (Serra *et al*., 2022). Auxin and vascular cambium initiation are critical signals for the establishment and maintenance of the cork cambium, and the transcription factors WOX4 and BP/KNAT1 act downstream of auxin, regulating cork cambium proliferation (Xiao *et al*., 2020) (Fig. 6). Suberin and lignin depositions in the cork are important for barrier function and cork cell integrity, as plants with reduced suberin and lignin levels have collapsed cork cells and are more susceptible to salt stress (Andersen *et al*., 2021). Still, how cork cells differentiate from the stem cells of the cork cambium and become highly suberized and lignified is largely unknown.

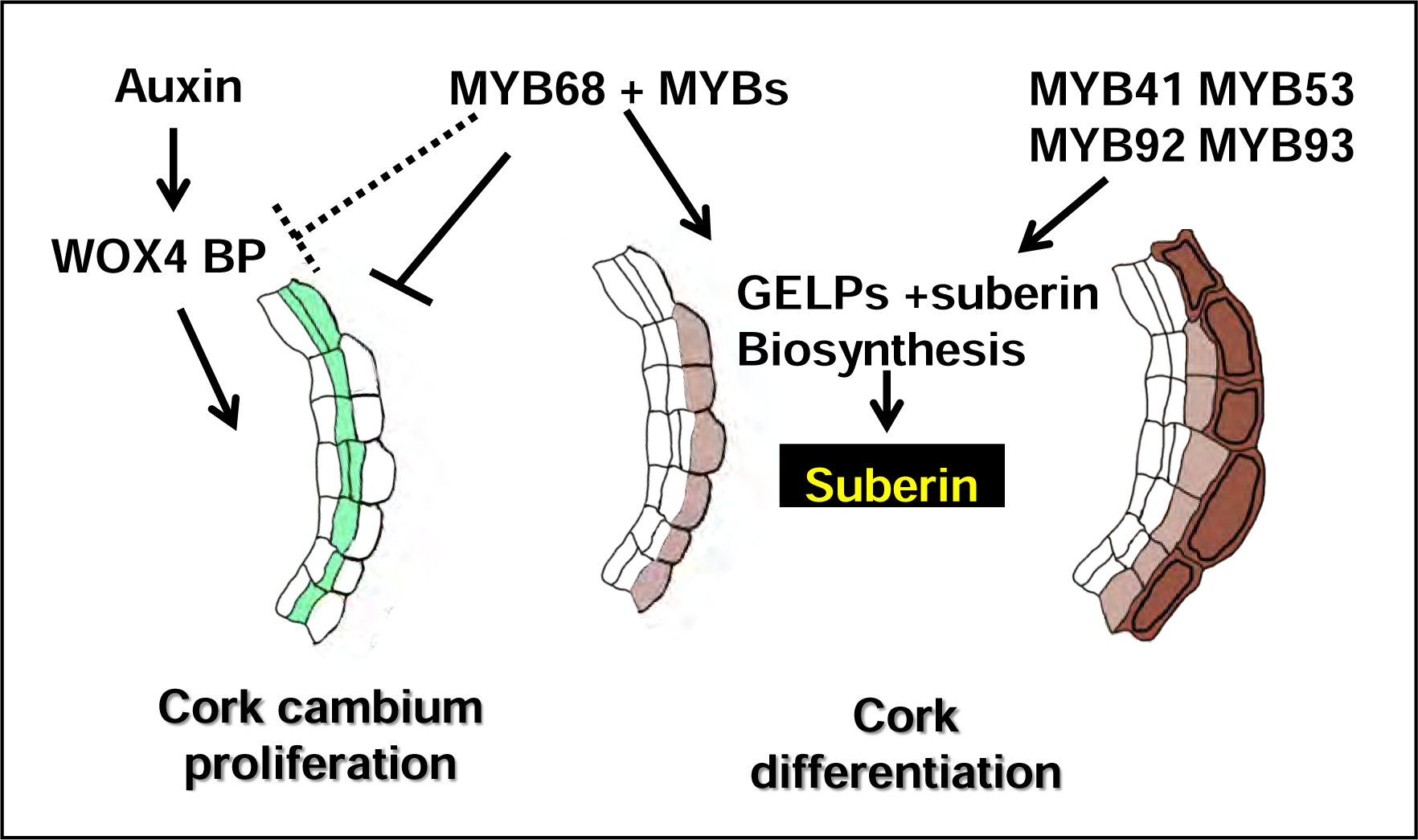

Our findings reveal that MYB68 and other members of the S14 subclade orchestrate cork differentiation by repressing cork cambium proliferation and promoting suberin deposition in cork cells (Fig. 6). MYB68 regulates suberin deposition at many levels: it promotes the expression of other regulators, such as MYB53, known to promote suberin deposition (Shukla *et al*., 2021) and enzymes essential for synthesizing aliphatic suberin components and suberin polymerization in the apoplast such as GDSL lipases/GELPs. Ursache and colleagues have recently shown that five GELPs are required for suberin deposition in the endodermis (Ursache *et al*., 2021). Interestingly, our genetic analysis revealed that the same GELPs are essential for suberin accumulation in the cork and are the targets of MYB68. Surprisingly, the expression of a single GELP was sufficient to restore the suberin phenotype of *myb68 myb84* double mutants, highlighting redundancy in the family and their importance in suberin deposition. In contrast to MYB36, MYB68 is able to trigger the expression of the suberin network and induce extra accumulation of suberin in the cork, revealing specificity among closely related transcription factors. However, we demonstrated that other MYBs that belong to different subclades and promote suberin in other developmental contexts, such as MYB41, MYB53, MYB92, and MYB93, also act in the cork, implying that suberin deposition is a robust process controlled by a complex network of transcription factors.

Unique to the MYB36 subclade is the role of supporting differentiation by repressing cell proliferation. In fact, only loss of function of MYB68 in combination with MYB84, MYB37, and MYB38 results in extra undifferentiated periderm layers and, in general, more peridermal cells. In addition, overexpression of MYB68 in the periderm triggers the exhaustion of the cork cambium and the production of more cork differentiated layers, a trait of agroeconomic interest as extra cork layers correlate with resistance to common scab disease in potato tuber and periderm thickness to wildfire survival in trees (Graves *et al*., 2014; Thangavel *et al*., 2016). The ability to regulate cell proliferation is not restricted to the context of periderm development; it encompasses other developmental processes. For instance, *myb36* mutants are characterized by extra ground tissue layers at the root tip (Kamiya *et al*., 2015; Liberman *et al*., 2015), MYB37 regulates leaf axil stem cell niche (Keller *et al*., 2006; Muller *et al*., 2006), while MYB87 has been shown to restrain vascular cambium activity (Zhang *et al*., 2019). Interestingly, ectopic expression of MYB68 in the vascular cambium mimics MYB87 overexpression, leading to consumption of the vascular cambium and extra xylem vessels. This suggests that MYB68 also has the ability to repress stem cell differentiation in other growth processes and that distinct expression patterns dictate which MYB controls stem proliferation in a specific developmental program.

In the cork cambium, MYB68 represses proliferation in parallel to or downstream of auxin, which is required for cork cambium maintenance and initiation, as interfering with auxin accumulation does not impede the formation of a thick periderm in *myb qudr.* mutants. However, WOX4 and BP genetically interact with MYB68 and MYB84 as they are required for the extra periderm layers that are produced by the cork cambium of *myb68 myb84* double knock-outs. Interestingly, MYB68 does not seem to regulate the expression of WOX4 and BP directly. Possible explanations include the idea that MYBs interact at the protein level with regulators of WOX4 and BP or that MYB68 and MYB84 bind to WOX4 and BP, interfering with their normal transcriptional outputs (FIG 6). This is a likely scenario as MYB transcription factors are known to form homodimers and heterodimers and to interact with other family of transcription factors to regulate developmental processes such as stomata formation and trichome differentiation (Pattanaik *et al*., 2014; Payne *et al*., 2000; Zhao *et al*., 2008).

In conclusion, our work illustrates the importance of MYB transcription factors in orchestrating the different steps of cork differentiation, highlighting both subclade-specific and unspecific functions. The notion that orthologs of MYB68 are expressed in cork cells of the stems of several trees and upon wounding in potato tubers goes along with the idea that there are conserved mechanisms between root, stem, and wound-induced cork differentiation. Supporting even a broader role for MYB68, it has been recently proposed to regulate suberin deposition in the endodermis and during apple fruit russeting (Xu *et al*., 2022; Xu *et al*., 2023). Finally, our work provides novel targets to engineer and manipulate barrier properties, such as the number of barrier cell layers and polymer deposition. Moreover, it will pave the way to dissect how the different barrier traits affect plant performance during development and under specific stress conditions.

## Supporting information

Supplemental Fiigures and Tables

## Acknowledgments

This work was supported by the Deutsche Forschungsgemeinschaft (DFG) with grant RA2590/4-1 and SFB1101 project B10 to LR. The LSM880 and the Arabidopsis walk-in chamber used in this study were acquired via DFG Major Research Instrumentation grants (INST 37/965-1 FUGG and INST 37/819-1 FUGG). TGA thanks the Sofja Kovalavskaya program at the Alexander von Humboldt Foundation as well as the Max Planck Society for funding. We thank Marie Barberon, Rochus Benni Franke, Niko Geldner, and Robertas Ursache for sharing seeds.

## Materials and Methods

### Plant Material and Growth

Unless stated otherwise, plants were grown in vitro in 1/2MS media at 22°C in continuous light. All Arabidopsis thaliana lines are in the Columbia background. Details of mutant alleles and transgenic lines are described in Table S1-S2. For dexamethasone (DEX) induction (Sigma, Cat# D1756), plants were initially grown and then, after either 9 or 14 days, transferred to 1/2MS media supplemented with 10μM DEX. For N-1-naphthylphthalamic acid (NPA) treatments (Duchefa Cat# N0926), plants were grown for 9 days on 1/2MS and then transferred to 1/2MS media supplemented with 10μM NPA. For salt stress experiments, plants were grown for 12 days on 1/2MS and then transferred to four-well split plates supplemented with 100 mM NaCl in one of the chambers. For all pharmacological treatments, unless specified otherwise, plants were treated for 5 days. For the qPCR experiments, 17-day-old plants were treated for 24 hours on 1/2MS media supplemented with 10μM DEX.

### Histological Techniques

For the analysis of root suberin, Fluorol Yellow (Santa Cruz Cat# sc-215052) was employed. The whole roots (for assessing root suberin coverage) or root periderm regions (for confocal suberin analysis) were first collected and placed in water. Subsequently, they were incubated in a solution of Fluorol Yellow 088 (0.01% dissolved in lactic acid) in the dark for 30 minutes at 70°C. The stained material was subsequently washed twice with water and incubated in a solution of aniline blue (0.5% in water) for counterstaining. After staining, plant material was mounted with 10% glycerol on glass slides for analysis using epifluorescence or confocal microscopy. As for plastic sections, the procedure remained the same, except in this case, Fluorol Yellow was dissolved in water, and the FY incubation was carried out at room temperature in the dark.

For periderm lignin studies, roots were collected in water and then stained in a solution of Basic Fuchsin (Sigma, 857343-100G) (0.5% in water) for 5 minutes. Afterward, they were washed three times with water, mounted with water on glass slides, and observed under the microscope.

For periderm anatomy studies, 1 cm root samples from the root junction were collected in water. Thin plastic cross-sections were obtained following the protocol described in (de Reuille and Ragni, 2017) using Technovit 7100 (Heraeus Kulzer Cat# 64709003). Plastic sections (5-7 µm) were stained with 0.1% toluidine blue and imaged with a Zeiss Axio M2 imager microscope.

For gene expression analysis, we employed two approaches: vibratome cross-sections and whole-mounts. First, in the case of vibratome cross-sections, 1 cm-root samples (0.5 cm below the root junction) were collected in water and then fixed using a 4% PFA solution (in 1xPBS) following the protocol described in (Singh *et al*., 2022; Ursache *et al*., 2018). The fixation was performed under vacuum for 1-2 hours, and then the root parts were rinsed twice for 1 minute in 1xPBS and embedded in 6% agarose blocks. Blocks were sectioned with a vibratome (Leica VT 1000). Approximately 20 vibratome cross-sections (50-100µm) were collected in water and then transferred to a clearing solution (ClearSee (Ursache *et al*., 2018)). Staining of vibratome cross-sections was prepared following the protocol described in (Singh *et al*., 2022; Ursache *et al*., 2018). Briefly, for Calcoflour White staining, vibratome cross-sections were stained with a 0.05% Calcoflour White solution (in ClearSee) for 5 minutes, while for Direct Red staining, vibratome sections were incubated for 30 minutes in a 0.05% Direct Red solution (in ClearSee). In both cases, the cross-sections were subsequently washed three times for 5 minutes each with the clearing solution and mounted on a glass slide using ClearSee. For whole-mount imaging, root segments were directly mounted into a solution of Propidium Iodide (PI, 10-20 mg/ml) or water for subsequent analysis.

### Confocal Laser Scanning Microscopy Imaging

Whole mount roots, roots vibratome sections, plastic cross-sections, or cleared cross-sections were imaged with a Zeiss LSM880 confocal microscope with the following settings: cork autofluorescence (ex. 405 nm; em. 420-460 nm), Basic Fuchsin (ex.566 nm; em.570-630 nm). FY (ex.488nm, em. 490-540 nm), GFP (488nm, 490 -510 nm), Venus and Citrine: (ex. 514 nm; em. 520–540 nm) and Calcofluor white (ex.475 nm, em 405-425nm). 3D reconstructions and orthogonal views of a Z stack were obtained using the ZEN Black software.

### Suberin quantification

For the determination of suberin content in the periderm, GC-MS (Gas Chromatography-Mass Spectrometry) was employed. Sample collection and the suberin extraction were performed following the protocol described in (Andersen *et al*., 2021). In brief, root sections of approximately 2 cm from the root junction were collected from 19-day-old plants. Most lateral roots were removed from the samples. Enzymatic treatment was utilized for cell wall degradation, followed by Soxhlet extraction with a chloroform: methanol mixture (1:1; v/v) for 2 days to remove unbound lipids. After the Soxhlet extraction, the samples were dried and weighed. Dried samples containing suberin were depolymerized using a 10% BF3/MeOH-based procedure. Before GC-MS analysis, samples were concentrated using N2 and derivatized with 20 μL of BSTFA (bis-(N,O-trimethylsilyl)-tri-fluoroacetamide, Macherey-Nagel, Germany) and 20 μL Pyridin for 40 min at 70°C. To quantify the suberin monomers detected by GC-MS (μg/mg dried sample), 25 μl of C32 alkane internal standard (13,5 mg/50 ml) were added to each sample.

### Image Analyses

In periderm anatomy studies, plastic cross-sections were used to quantify the number of periderm and cork cells. Initially, bright-field cross-section images were used to determine the total count of periderm cells. Subsequently, fluorescent microscopy was applied to the same plastic cross-sections to ascertain the number of cork cells. Cross-sections were exposed to UV excitation through a 365 nm DAPI filter to observe the intrinsic autofluorescence of mature cork cells. The total number of periderm cells and the ratio of cork cells to the total number of periderm cells are presented in the results. Three independent experiments were conducted, and only one was presented. The cork ratio: cork length in cm / total root length in cm was measured as described in (Wunderling *et al*., 2018).

### Molecular Cloning

The Green Gate cloning system, described in (Lampropoulos *et al*., 2013), was used to create the majority of constructs used in this study (see Table S3-S4). Promoters were amplified from genomic DNA and the coding sequences from root cDNA. Primers used for the cloning are listed in Table S5. All modules and vectors used in this study are described in Table S3. The module assembly strategy is described in Table S4. The promoter of MYB87 was cloned using Gateway technology in pDNOR207 and then introduced into a final destination vector containing NLS-GFP. The promoter of MYB38 (1638bp) was infusion cloned into a L1R4 GateWay vector in a similar manner as described in (Andersen *et al*., 2021) All final constructs were transformed into Agrobacterium tumefaciens GV3101 together with pSoup for plant transformation via floral dipping or transactivation assays.

#### Genome editing

The cloning of plasmids used for genome editing of MYB87 and MYB68 via CRISPR-Cas9 was conducted following the procedure outlined in (Fauser *et al*., 2014). The following gRNA was used for MYB87: ATTGACCGTGCTGCGACAAGATGG and for MYB68: ATCGAGAATAGTGGCACAGG. MYB87 and MYB68 sgRNA primers are listed in Table S5. In brief, oligos were annealed and subsequently ligated into BbsI-linearized Gateway-entry plasmid pEn-Chimera. Finally, MYB87 and MYB68 sgRNA were introduced into a gateway destination vector for plant transformation harboring the Cas9 enzyme. The final constructs were introduced into Col-0 plants using floral dipping. Positive Fast-red-T1 seeds were selected, cultivated in soil, and genotyped. T2 seeds from edited T1 plants that do not carry the CAS9 vector were selected (no red fluorescent seeds), propagated, and genotyped to obtain homozygous MYB87 and MYB68 edited plants. For genotyping, the region of interest was amplified using the primers described in Table S5 and analyzed by sequencing.

### Sample preparation for RNA-seq and qPCR experiments

RNA was extracted from root sections of approximately 2 cm from the root junction. Lateral roots were excised, and the samples were promptly collected in a tube submerged in liquid nitrogen to preserve RNA integrity. To create one single RNA sample, between 40 and 50 root sections were pooled together. The RNA extraction was done using the Universal RNA Purification Kit (Roboklon, E3598-02) according to the manufacturer’s protocol.

### RNAseq data processing and analysis

RNA sequencing (paired-end 150bp) was performed at Novogene. Raw data sequences will be available after publishing. Data processing and data analysis were performed using the Galaxy platform (https://usegalaxy.eu/)(Afgan *et al*., 2018). Adaptors were removed with Trim Galore using default parameters, and read quality was assessed with FastQC. Reads were aligned to the TAIR10 genome using HISAT2 with default parameters. Reads were counted using featureCounts, and differential gene expression analysis was performed using DESeq2 and can be queried in DataS1.

### qPCR analysis

C-DNA was synthesized using AMV Reverse Transcriptase Native (Roboklon, E1372-01). For qPCR MESA blue (Eurogentec, RT-SYS2X-03-+NRWOUB) was used, and the reactions were performed in a CFX96 Real-Time System machine (BIO-RAD). Primers used for qPCR are listed in Table S5. The relative expression was calculated using CFX Maestro software (BIO-RAD), and the samples were normalized against EF1. qPCR experiments were repeated at least 3 times, and one experiment was shown.

### Transient Transactivation Studies in Nicotiana benthamiana Leaves

Using the GreenGate cloning system, we assembled two types of constructs for expression in plants. First, vectors containing the 35S promoter and our transcription factors of interest (MYB36, MYB68, or nothing as control) tagged with mCherry at the C-terminus (see Table S3-S5) were obtained. Second, we generated constructs with the fungal luciferase reporter (*N. nambi* luciferase, nnLuz) under the control of our promoters of interest (see Table S3-S5). Subsequently, these vectors were then transformed into GV3101 Agrobacterium strain. We prepared liquid cultures and grew them until reaching an OD 600 of 0.8. Then, we co-infiltrated 3-week-old *N. benthamiana* leaves with 3 constructs: 1) the vector containing the transcription factor of interest, 2) the vector containing the promoter of interest and the luciferase reporter, and 3) the P336-FBP_11 vector (Available from AddGene (#139702)), which contains the biosynthesis pathway for luciferin production and recycling ((Khakhar *et al*., 2020). For co-infiltration, the Agrobacterium culture of each vector was diluted to an OD of 0.250, resulting in a total OD of 0.750 for each leaf. The mixture was washed with water twice and then resuspended in an infiltration medium composed of 10mM MgCl2, 1mM MES, and 100μM Acetosyringone. The resuspended solution was incubated for 1 hour and subsequently infiltrated into tobacco leaves. At 3 day-post-infiltration (dpi), images of luminescence of the infiltrated leaves were acquired using a CCD camera (Amersham 600 gel imager) following a 3-minute exposure.

### Quantitative Analysis

No statistical method was employed to predetermine the sample size, and the experiments were not randomized. IBM SPSS Statistics version 24-25-26 (IBM) was used to make statistical analysis. The number of periderm and cork cells, the root length, and the root suberin coverage were determined using ImageJ / Fiji5. The root length and root suberin coverage were measured according to (Andersen *et al*., 2021). For the suberin root coverage quantification in old roots that comprised the periderm, the statistic was performed separately for each root zone.

For multiple sample comparisons, a one-way ANOVA with Tukeys’s post hoc test (equal variance not assumed) was performed unless otherwise stated. For GC-MS and periderm experiments, statistical analyses were performed using IBM SPSS Statistics version 25 (IBM). First, the datasets were tested for the homogeneity of variances using Levene’s Test. The significant differences between the two datasets were calculated using Welch’s two-tailed t-test in case of a non-homogeneous variance or a Student’s two-tailed t-test if the variance was homogeneous. For the detailed suberin quantification of each compound/category of compounds via GC-MS, the statistic was performed separately for each compound/category.

## Supplemental Figures

**Figure S1** (A) Orthogonal view of z stacks showing the expression of MYB68 (*pMYB68:NLS-3xGFP W131Y)*, MYB84 (*MYB84:NLS-3xGFP W131Y*), MYB87 (*pMYB87:NLS-GFP*), MYB37 (*pMYB37:NLS-GFP*), MYB38 (*pMYB38:NLS-Scarlet*) and MYB36 (*pMYB36g:NLS-GFP myb36*) at two periderm developmental stages (14-day-old roots). W131Y (a ubiquitous plasma membrane marker (*UBQ10:eYFP-NPSN12*)), PI (Propidium Iodide), or the intrinsic autofluorescence of cork cells, was employed to outlay cells. The white arrows indicate the expression in the cork. (B) Confocal microscopy images of vibratome cross-sections of 19-day-old roots showing the expression of *MYB37*, *MYB38*, *MYB36*, and *MYB87* in the periderm using the same lines used in (A). The white arrowheads indicate the cork cells. (C) Orthogonal view of z stacks showing the expression of *MYB68* and *MYB84* together with cork (*GPAT5:mCitrine-SYP122)* differentiating cork/cork cambium (*PER15:mCherry-SYP122*), and cork cambium/phelloderm (*WOX4:YFPer*) reporters. (D and E) Representative images of the relative intensity of FY signal in the cork. (D) WT, *myb68*, *myb84*, *myb87* and *myb37 myb38 myb84*. (E), WT and *myb36.* White scale bars: 20µm.

**Figure S2** (A) Representative images of relative intensities of Fluorol Yellow(FY) fluorescent signal in the periderm of 14-day-old WT, *myb68 myb87*, *myb68 myb84*, and *myb quadr*. roots. (B-D) Quantification of percentage coverage of developing and mature cork of 19-day-old WT, *myb36*, *myb68*, *myb84*, *myb87*, *myb68 myb87*, *myb68 myb84*, *myb37 myb38 myb84* and *myb quadr*. roots. (E) Root length (cm) of 14-day-old WT, *myb68 myb84,* and *myb quadr*. roots. (F) Quantification of the cork ratio of 14-day-old WT, *myb68 myb84,* and *myb quadr*. roots. Data were obtained as described in the methods section. (G) Relative quantification of developing and mature cork suberin regions of 19-day-old WT and pGPAT5:MYB68-GR *myb68 myb84* roots. 9-day-old plants were treated for 5 days with mock or 10μM DEX. (H) Representative images of the relative intensities of FY fluorescent signal in the periderm of 14-day-old WT and *pGPAT5:MYB68-GR myb68myb84* roots. 9-day-old plants were treated for 5 days with mock or 10μM DEX. (I) Representative images of the relative intensities of basic fuchsin fluorescent signal of 14-day-old WT, *myb68myb84*, and *myb quadr* roots. In (C, E-F,G) one-way ANOVA (95% confidence interval [CI], post hoc: Tukey HSD, (n = 10-14). In (B,D) Two-tailed Students T-Test ( *: P<0.05, n:15-20) White scale bars: 20µm.

**Figure S3** (A) Set of differentially expressed genes in WT vs. *myb68 myb84,* and Cork enriched genes *vs.* all tissues (data from (Leal *et al*., 2022)). The log 2 Fold Changes (FC) and adjusted P value are highlighted by a color code. (B-E). Transactivation studies in *N. benthamiana* leaves using luciferase reporting system. Upper panels show the luminescence intensities of leaves co-infiltrated with an effector construct (*p35S:mCherry*, *p35S:MYB68-mCherry*, *p35S:MYB84-mCherry*), a reporter construct (promoter of interest with the luciferase reporter: *pFAR4:LUX*, *pGPAT5:LUX*, *pMYB53:LUX* and *pCASP2:LUX*), and the FBP11 construct (encoding the fungal bioluminescent pathway). Lower panels show the RGB images of the same leaves. Black scale bars: 1cm.

**Figure S4** (A) Cross-sections (plastic embedding) of the uppermost part of 14-day-old WT and *myb37 my38 myb84.* (B-C) Quantification of the ratio of the number of cork cells/number of periderm cells and the total number of periderm cells of the experiment presented in (A). (D) Cross-sections of the uppermost part of 14-day-old WT, *myb87*, *myb68*, *myb84*, *myb68myb87*, *myb68myb8*4, *myb quadr*. Black double-headed arrows show the periderm area. (E-F) Quantification of the ratio of the number of cork cells/number of periderm cells and the total number of periderm cells of the experiment presented in (D). (G) Cross-sections of 14-day-old WT*, myb68myb84*, and *myb quadr*. roots. It shows autofluorescence in response to UV radiation excitation of cork cells. Red arrows show the developing cork characterized by the lack of fluorescence. (H-I) Fluorol Yellow (FY.) Staining and Cork suberin coverage of WT and *ralph horst* roots. (H) Relative quantification of developing and mature cork suberin regions of 19-day-old plants. (I) Relative intensities of Fluorol Yellow fluorescent signal in the periderm of 14-day-old plants. (J) Cross-sections of the uppermost part of 14-day-old *pPXY:MYB68-GR* and *pPXY:MYB87-GR*. Two independent lines of *pPXY:MYB87-GR* are shown. 9-day-old plants were treated for 5 days with mock or 10μM DEX. In (B, C, and H) Two-tailed Students T-Test ( *: P<0.05, (B and C) n:15-20, (H) n:10-15). In (E-F) One-way ANOVA (95% confidence interval [CI], post hoc: Tukey HSD, (n = 15-20). Black double-headed arrows show the periderm area. Scale bars (black and white) in cross sections 20µm.

## Notes

### Competing Interest Statement

The authors have declared no competing interest.

