## Supplemental Fiigures and Tables for "MYB68 orchestrates cork differentiation by regulating stem cell proliferation and suberin deposition"

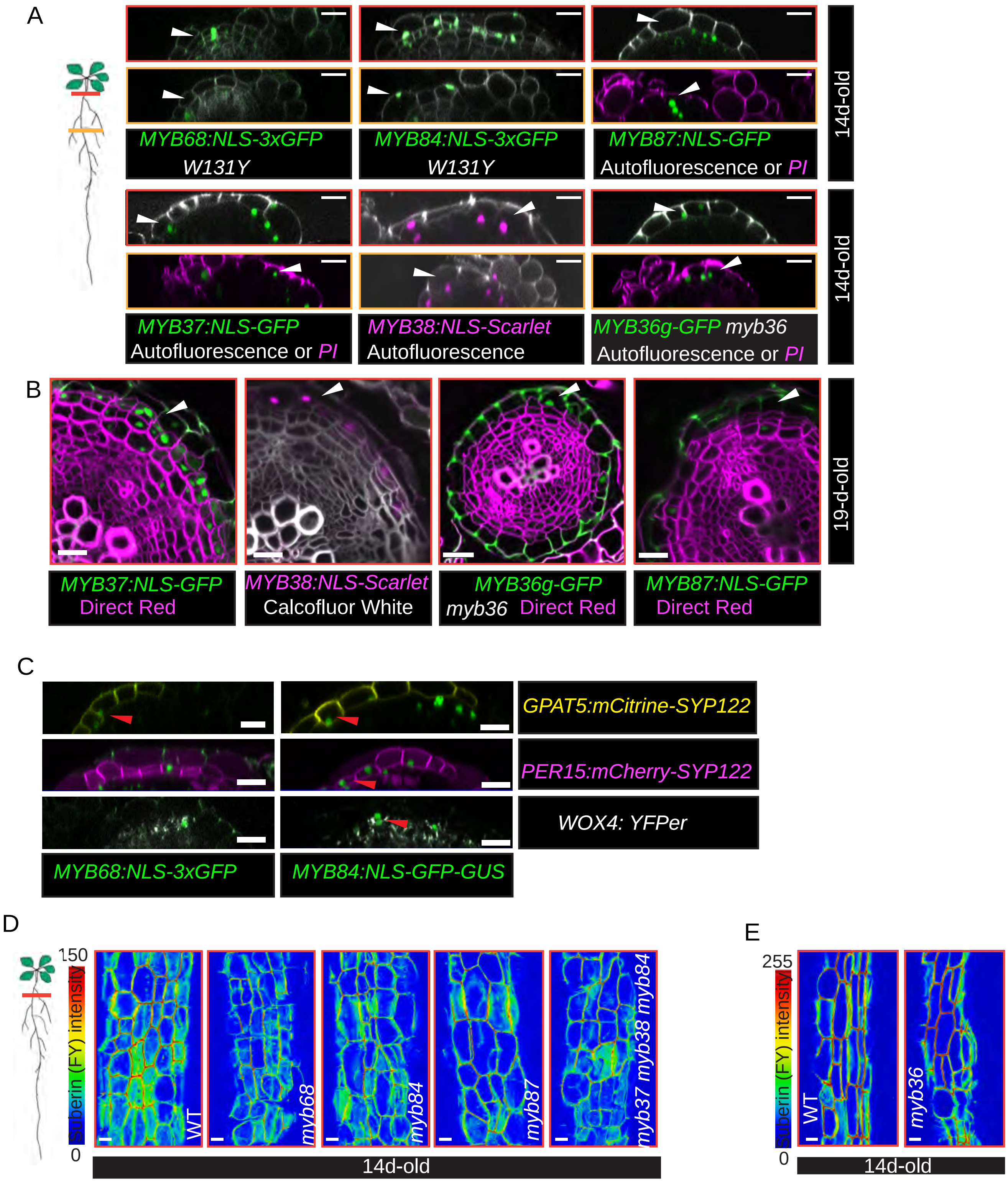

A
MYB68:NLS-3xGFP
W131Y
MYB84:NLS-3xGFP
W131Y
MYB87:NLS-GFP
Autofluorescence or PI
14d-old
MYB37:NLS-GFP
Autofluorescence or PI
MYB38:NLS-Scarlet
Autofluorescence
MYB36g-GFP myb36
Autofluorescence or PI
14d-old
B
MYB37:NLS-GFP
Direct Red
MYB38:NLS-Scarlet
Calcofluor White
MYB36g-GFP
myb36 Direct Red
MYB87:NLS-GFP
Direct Red
19-d-old
C
GPAT5:mCitrine-SYP122
PER15:mCherry-SYP122
WOX4: YFPer
MYB68:NLS-3xGFP
MYB84:NLS-GFP-GUS
D
150
Suberin (FY) intensity
0
WT
myb68
myb84
myb87
myb37 myb38 myb84
14d-old
E
255
Suberin (FY) intensity
0
WT
myb36
14d-old

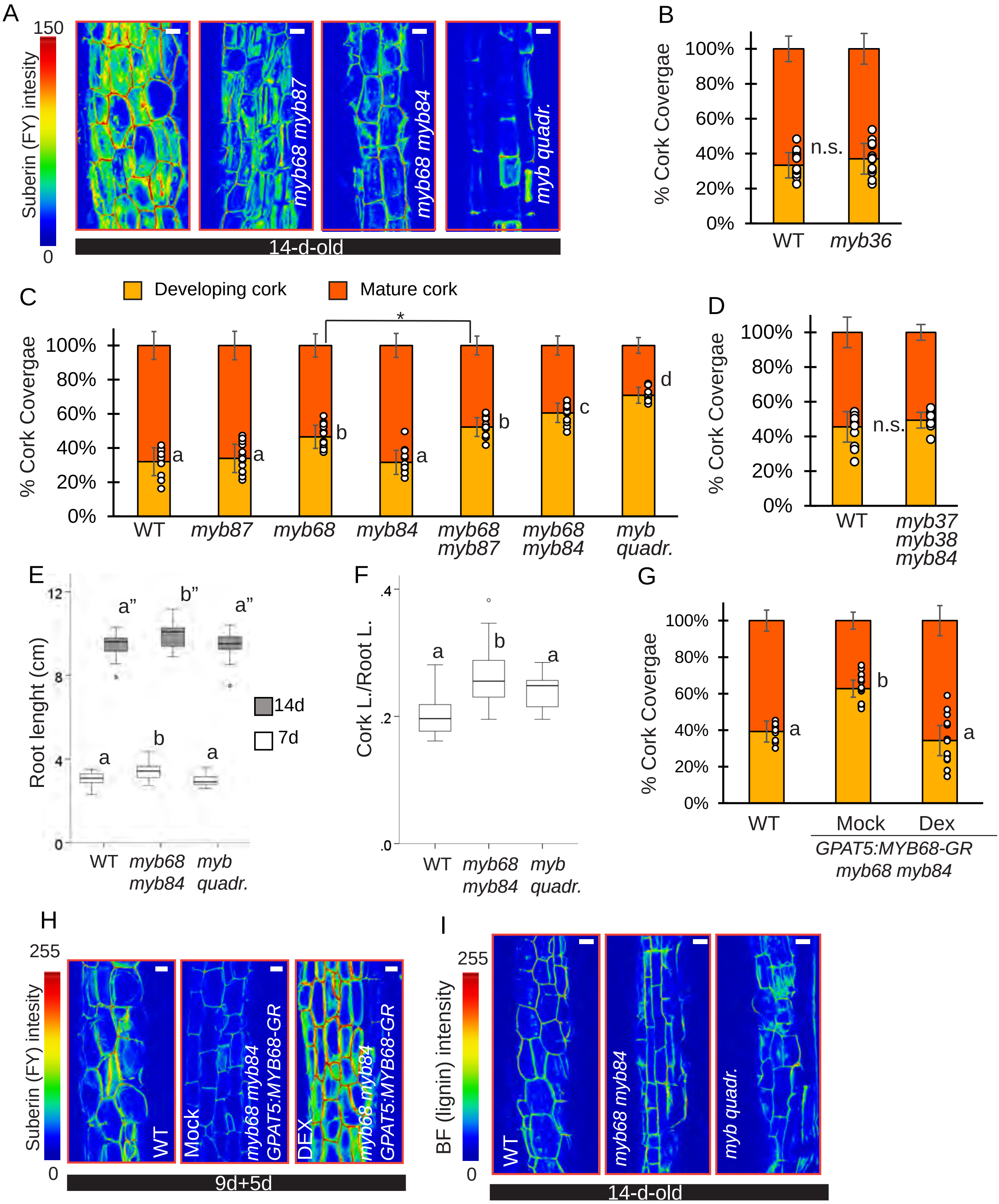
A
150
Suberin (FY) intesity
0
myb68 myb87
myb68 myb84
myb quadr.
14-d-old
B
% Cork Covergae
100%
80%
60%
40%
20%
0%
n.s.
WT
myb36
C
Developing cork
Mature cork
*
a
a
b
a
b
c
d
WT
myb87
myb68
myb84
myb68 myb87
myb68 myb84
myb quadr.
D
n.s.
myb37 myb38 myb84
E
Root lenght (cm)
12
8
4
0
a"
b"
a"
a
b
a
14d
7d
WT
myb68 myb84
myb quadr.
F
Cork L./Root L.
.4
.2
.0
a
b
a
G
a
b
a
Mock
Dex
GPAT5:MYB68-GR myb68 myb84
H
255
Suberin (FY) intesity
WT
Mock
myb68 myb84 GPAT5:MYB68-GR
DEX
9d+5d
I
255
BF (lignin) intensity
WT
myb68 myb84
myb quadr.
14-d-old

A

| Gene Name | Gene Number | WT *vs* *myb68 myb84* log2 (FC) | WT *vs* *myb68 myb84* Padj | Cork *vs* All tissues log2 (FC) | Cork *vs* All tissues Padj |
|---|---|---|---|---|---|
| **KCS2** | AT1G04220 | -0.2096 | 3.2E-01 | 2.77535 | 2.8E-108 |
| **FAR1** | AT5G22500 | -0.46373 | 5.2E-08 | 4.02325 | 6.2E-217 |
| **FAR4** | AT3G44540 | -0.55878 | 4.0E-06 | 2.91422 | 3.4E-127 |
| **FAR5** | AT3G44550 | -0.5692 | 1.5E-05 | 3.01793 | 3.9E-141 |
| **HORST** | AT5G58860 | -0.52038 | 1.3E-08 | 2.39351 | 2.9E-68 |
| **KCS20** | AT5G43760 | -0.5495 | 7.0E-08 | 0.71423 | 1.8E-07 |
| **GPAT5** | AT3G11430 | -0.4574 | 8.3E-04 | 2.09063 | 4.5E-41 |
| **RALPH** | AT5G23190 | -0.35883 | 7.2E-03 | 1.96528 | 1.2E-41 |
| **FACT** | AT5G63560 | -0.44276 | 2.2E-07 | 2.96926 | 3.6E-172 |
| **GELP8** | AT1G28590 | -1.68939 | 7.3E-21 | 2.02028 | 1.5E-10 |
| **GELP49** | AT2G19050 | -1.07764 | 7.8E-07 | -0.35252 | NA |
| **GELP16** | AT1G29670 | -1.02618 | 9.3E-09 | -1.03629 | 4.3E-03 |
| **GELP20** | AT1G53940 | -0.87642 | 1.8E-08 | -0.90834 | 2.7E-01 |
| **GELP100** | AT5G45670 | -0.71291 | 8.6E-05 | -0.53723 | 3.8E-01 |
| **GELP102** | AT5G45950 | -0.61461 | 1.2E-02 | -1.24362 | 5.7E-02 |
| **GELP69** | AT3G27950 | -0.59009 | 1.4E-02 | 1.42546 | 8.9E-02 |
| **GELP67** | AT3G16370 | -0.58804 | 1.2E-05 | -2.21072 | 1.5E-37 |
| **GELP103** | AT5G45960 | -0.57349 | 3.4E-02 | 1.43564 | 4.4E-02 |
| **GELP63** | AT3G14210 | -0.56307 | 2.5E-02 | 1.16349 | 1.9E-01 |
| **GELP96** | AT5G37690 | -0.54612 | 1.0E-06 | 2.09176 | 2.0E-65 |
| **GELP51** | AT2G23540 | -0.48493 | 4.0E-08 | 1.82549 | 2.6E-49 |
| **GELP41** | AT1G75900 | -0.4468 | 1.2E-01 | -1.36994 | 1.0E-03 |
| **GELP73** | AT3G50400 | -0.43842 | 6.8E-03 | 1.28259 | 9.5E-15 |
| **GELP38** | AT1G74460 | -0.43497 | 1.5E-04 | 1.73705 | 1.6E-39 |
| **GELP22** | AT1G54000 | -0.26539 | 4.7E-02 | 0.29083 | 1.4E-01 |
| **MYB41** | AT4G28110 | 0.16726 | 7.3E-01 | 0.98029 | 1.7E-02 |
| **MYB93** | AT1G34670 | -0.05159 | 8.6E-01 | 1.59116 | 4.0E-31 |
| **MYB53** | AT5G65230 | -0.38532 | 1.6E-01 | 1.24628 | 1.1E-09 |
| **MYB92** | AT5G10280 | 0.17894 | 2.9E-01 | 1.89528 | 2.9E-29 |
| **MYB39** | AT4G17785 | 0.01933 | 9.7E-01 | 1.92438 | 3.2E-06 |
| **MYB107** | AT3G02940 | 0.23685 | 0.4895 | 1.57173 | 2.8E-21 |
| **MYB74** | AT4G05100 | 0.80997 | 8E-06 | 2.39236 | 1.1E-40 |
| **MYB9** | AT5G16770 | 0.03333 | 0.956 | 1.35888 | 1.1E-14 |
| **MYB102** | AT4G21440 | 0.58926 | 2.6E-02 | 3.20486 | 3.5E-27 |
| **ANAC46** | AT3G04060 | -0.11147 | 5.9E-01 | 0.42101 | 1.9E-01 |
| **WOX4** | AT1G46480 | 0.04027 | 9.2E-01 | | |
| **BP** | AT4G08150 | 0.47762 | 6.5E-05 | | |

**-2 log2 (FC)+2**

**-4 log2 (FC)+4**

**Padj < 0.01**

**Padj ≥ 0.01**

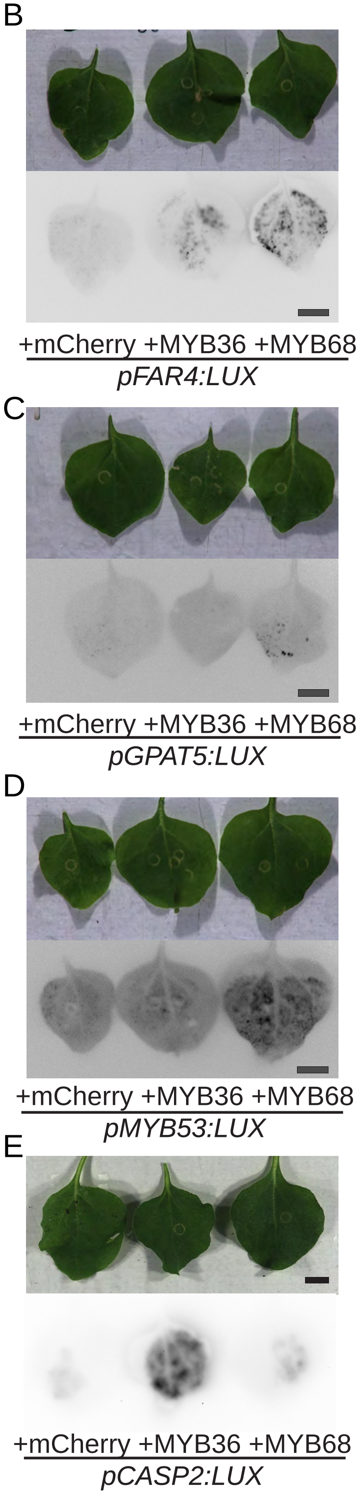

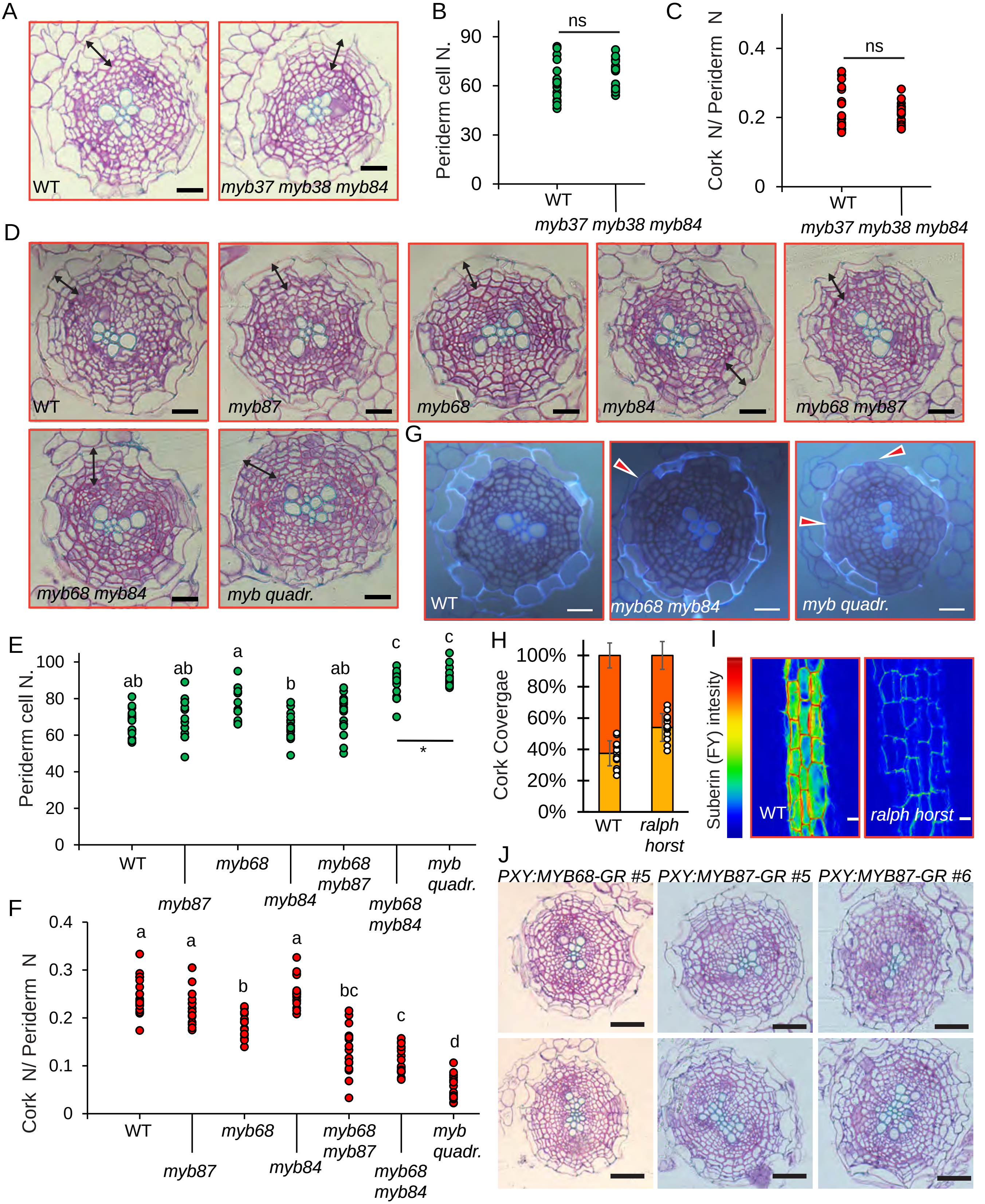

A
WT
myb37 myb38 myb84
B
ns
Periderm cell N.
90
60
30
0
WT
myb37 myb38 myb84
C
ns
Cork N/ Periderm N
0.4
0.2
0
WT
myb37 myb38 myb84
D
WT
myb87
myb68
myb84
myb68 myb87
myb68 myb84
myb quadr.
G
WT
myb68 myb84
myb quadr.
E
Periderm cell N.
100
80
60
40
20
0
ab
ab
a
b
ab
c
c
*
WT
myb87
myb68
myb84
myb68 myb87
myb68 myb84
myb quadr.
H
Cork Covergae
100%
80%
60%
40%
20%
0%
WT
ralph horst
I
Suberin (FY) intesity
WT
ralph horst
F
Cork N/ Periderm N
0.4
0.3
0.2
0.1
0
a
a
b
a
bc
c
d
WT
myb87
myb68
myb84
myb68 myb87
myb68 myb84
myb quadr.
J
PXY:MYB68-GR #5
PXY:MYB87-GR #5
PXY:MYB87-GR #6

**Table S1 Transgenic lines used in this study**

| **Arabidopsis Lines** | **Obtained from/by** | **Described in** |
|---|---|---|
| MYB84:NLS-3xGFP | Our work | (Wunderling et al. 2018) |
| MYB68:NLS-3xGFP | This study | This study |
| MYB84:NLS-3xGFP<br>W131Y (UBQ10::eYFP-NPSN12) | Our work | (Wunderling et al. 2018)<br>(Geldner et al. 2009) |
| MYB68:NLS-3xGFP<br>W131Y (UBQ10::eYFP-NPSN12) | This study | This study<br>(Geldner et al. 2009) |
| MYB87:NLS-GFP | This study | This study |
| MYB37:NLS-3xGFP | This study | This study |
| MYB38:NLS-3xScarlet | This study | This study |
| MYB36g-GFP in myb36-1 | N69051 | (Liberman et al. 2015) |
| MYB68:NLS-3XGFP<br>WOX4:ER-YFP | Crossing | This study<br>(Suer et al. 2011) |
| MYB68:NLS-3XGFP<br>GPAT5:mCitrine-SYP122 | Crossing | (Wunderling et al. 2018)<br>(Barberon et al. 2016) |
| MYB84:NLS-GFP-GUS<br>WOX4:ER-YFP | Crossing | (Wunderling et al. 2018)<br>(Suer et al. 2011) |
| MYB84:NLS-GFP-GUS<br>GPAT5:mCitrine-SYP122 | Crossing | (Wunderling et al. 2018)<br>(Barberon et al. 2016) |
| GPAT5:MYB68-GR in<br>MYB84:NLS-GFP GUS<br>GPAT5:mCitrine-SYP122<br>PER15:mCherry-SYP122 | This study | This study<br>(Wunderling et al. 2018)<br>(Barberon et al. 2016)<br>This study |
| GPAT5:MYB36-GR | This study | This study |
| GPAT5:MYB68-GR in<br>myb68-2-4 myb84-1 | This study | This study |
| PER15:MYB68-GR | This study | This study |
| PER15:MYB36-GR | This study | This study |
| GPAT5:GELP51 in myb68-2-4 myb84-1 | This study | This study |
| GELP22:NLS-3xmVenus | Robertas Ursache | (Ursache et al. 2021) |
| GELP38:NLS-3xmVenus | Robertas Ursache | (Ursache et al. 2021) |
| GELP49:NLS-3xmVenus | Robertas Ursache | (Ursache et al. 2021) |

| | | |
|---|---|---|
| GELP51:NLS-3xmVenus | Robertas Ursache | (Ursache et al. 2021) |
| GELP96:NLS-3xmVenus | Robertas Ursache | (Ursache et al. 2021) |
| MYB41:NLS-3xmVenus | Marie Barberon | (Shukla et al. 2021) |
| MYB93:NLS-3xmVenus | Marie Barberob | (Shukla et al. 2021) |
| MYB53:NLS-3xmVenus | Marie Barberon | (Shukla et al. 2021) |

**Table S2 Mutant alleles used in this study**

| **Name used in the paper** | **Allele** | **Atg** | **Type** | **Obtained from/by** | **Described in** |
|---|---|---|---|---|---|
| myb84 | myb84-1 | AT3G49690 | Salk_141918C | N653820 | (Feng et al. 2004) |
| myb68 | myb68-2-4 | AT5G65790 | CRISPR (extra C at position 100 of CDS that causes frame shift) | This study | This study |
| myb87 | myb87-3a | AT4G37780 | CRISPR (extra A after position 28 of CDS that causes frame shift) | This studyl | This study |
| myb36 | myb36-1 in MYB36::MYB36-GR | AT5G57620 | EMS | N69054 | (Liberman et al. 2015) |
| myb37<br>myb38<br>myb84 | rax1-3<br>rax2-1<br>rax3-1 | AT3G49690<br>AT5G23000<br>AT5G65790 | SALK_071748C<br>Wisconsin line;<br>rax3 En-1 | Sónia Gonçalves | (Muller et al. 2006) |
| myb68<br>myb84 | myb68-2-4<br>myb84-1 | AT3G49690<br>AT5G65790 | CRISPR<br>Salk_141918C | Crossing | This study |
| myb68<br>myb87 | myb68-2-4<br>myb87-3b | AT3G49690<br>AT5G65790 | CRISPR<br>CRISPR | This study | This study |
| myb quad. | rax1-3<br>rax2-1<br>rax3-1<br>myb68-2-4 | AT3G49690<br>AT5G23000<br>AT5G23000<br>AT5G65790 | SALK_071748C<br>Wisconsin line;<br>rax3 En-1<br>CRISPR | This study | This study |

| | | | | | |
|---|---|---|---|---|---|
| horst<br>ralph | horst-1<br>ralph-1 | AT5G58860<br>AT5G23190 | SALK_107454<br>SM.37066 | Rochus Franke | (Salas-González et al. 2021) |
| gelp q1 | gelp22-c1<br>gelp38-c3<br>gelp49-c1<br>gelp51-c1<br>gelp96-c1 | AT1G54000<br>AT1G74460<br>AT2G19050<br>AT2G23540<br>AT5G37690 | CRISPR<br>CRISPR<br>CRISPR<br>CRISPR<br>CRISPR | Robertas Ursache | (Ursache et al. 2021) |
| gelp q2 | gelp22-c1<br>gelp38-c3<br>gelp49-c1<br>gelp51-c1<br>gelp96-c1 | AT1G54000<br>AT1G74460<br>AT2G19050<br>AT2G23540<br>AT5G37690 | CRISPR<br>CRISPR<br>CRISPR<br>CRISPR<br>CRISPR | Robertas Ursache | (Ursache et al. 2021) |
| wox4 | wox4-1 | AT1G46480 | GK_462GO1 | Thomas Greb | (Suer et al. 2011) |
| bp | bp-9 | AT4G08150 | Transposon insertion | Pautot Veronique | (Mele et al. 2003) |
| myb68<br>myb84<br>bp | myb68-2-4<br>myb84-1<br>bp-9 | AT3G49690<br>AT5G65790<br>AT4G08150 | CRISPR<br>Salk_141918C<br>Transposon insertion | This study | This study |
| myb68<br>myb84<br>wox4 | myb68 2-4<br>myb84-1<br>wox4-1 | AT3G49690<br>AT5G65790<br>AT1G46480 | CRISPR<br>Salk_141918C<br>GK_462GO1 | This study | This study |
| myb41<br>myb53<br>myb92<br>myb93 | myb41_c2<br>myb53_c1<br>myb92_c1-<br>myb93_c1 | AT4G28110<br>AT5G65230<br>AT5G10280<br>AT1G34670 | CRISPR<br>CRISPR<br>CRISPR<br>CRISPR | Marie Barberon | (Shukla et al. 2021) |

Table S3 Greengate cloning mudles used in this study

| GG modules/vectors used in this study | Ref. | Obtained |
|---|---|---|
| pGG-A0 | (Lampropoulos et al. 2013) | Jan Lohmann |
| pGG-C0 | (Lampropoulos et al. 2013) | Jan Lohmann |
| pZ03 | (Lampropoulos et al. 2013) | Jan Lohmann |
| pGG-A-pGPAT5 | Baed on (Naseer et al. 2012) | This study |
| pGG-A-pFAR4 | Based on (Naseer et al. 2012) | This study |
| pGG-A-pCASP2 | Based on (Roppolo et al. 2011) | This study |
| pGG-A-pGELP8 | This study | This study |
| pGG-A-pGELP51 | Based on (Ursache et al. 2021) | This study |
| pGG-A-pGELP38 | Based on (Ursache et al. 2021) | This study |
| pGG-A-pPER15 | (Xiao et al. 2020) | Our group |
| pGG-A-pMYB68 | This study | This study |
| pGG-A-pMYB84 | (Wunderling et al. 2018) | Our group |
| pGG-A-pMYB37 | This study | This study |
| pGG-A-pMYB53 | This study | This study |
| pGG-A-p35S | (Lampropoulos et al. 2013) | Jan Lohmann |
| pGG-B03 (B-dummy) | (Lampropoulos et al. 2013) | Jan Lohmann |
| pGG-B05 (NLS) | (Lampropoulos et al. 2013) | Jan Lohmann |
| pGG-C25 (3xGFP) | (Lampropoulos et al. 2013) | Jan Lohmann |
| pGG-C-*MYB68* | This study | This study |
| pGG-C-*MYB36* | This study | This study |
| pGG-C-GELP51 | This study | This study |
| pGG-D-GR | (Ramakrishna et al. 2019) | Thomas Greb |
| pGG-D2 (D-dummy) | (Lampropoulos et al. 2013) | Jan Lohmann |
| pGG-E1 (pea RBCS terminator) | (Lampropoulos et al. 2013) | Jan Lohmann |
| pGG-F5 (Hygro R) | (Lampropoulos et al. 2013) | Jan Lohmann |
| pGG-F-FR (FASTRED) | (Vilches Barro et al. 2019) | Alexis Maizel |
| pGG-C-NnLUX | Based on (Khakhar et al. 2020) | This study |
| pGG-C-mCherry | (Lampropoulos et al. 2013) | Jan Lohmann |
| pGG-D-mCherry | (Lampropoulos et al. 2013) | Jan Lohmann |

**Table S4 GreenGate assembly of final constrcuct used in this study**

| **Final vector (in pZ03)** | A | B | C | D | E | F |
|---|---|---|---|---|---|---|
| *MYB68:NLS-3xGFP* | *pMYB68* | B5 | *C25* | D2 | E1 | F5 |
| *MYB37:NLS-3xGFP* | *pMYB37* | B5 | *C25* | D2 | E1 | F5 |
| *GPAT5:MYB68-GR* | *pGPAT5* | B3 | MYB68 | GR | E1 | FR |
| *GPAT5:MYB36-GR* | *pGPAT5* | B3 | MYB36 | GR | E1 | FR |
| *PER15:MYB68-GR* | *pPER15* | B3 | MYB68 | GR | E1 | FR |
| *GPAT5:GELP51* | *pGPAT5* | B3 | GELP51 | D2 | E1 | FR |
| *pGPAT5:LUX* | *pGPAT5* | B3 | NnLUX | D2 | E1 | FR |
| *pFAR4:LUX* | *pFAR4* | B3 | NnLUX | D2 | E1 | FR |
| *pCASP2:LUX* | *pCASP2* | B3 | NnLUX | D2 | E1 | FR |
| *pGELP38:LUX* | *pGELP36* | B3 | NnLUX | D2 | E1 | FR |
| *pGELP51:LUX* | *pGELP51* | B3 | NnLUX | D2 | E1 | FR |
| *pGELP8:LUX* | *pGELP8* | B3 | NnLUX | D2 | E1 | FR |
| *pMYB53:LUX* | *pMYB53* | B3 | NnLUX | D2 | E1 | FR |
| *35S:mCherry* | *p35S* | B3 | m-Cherry | D2 | E1 | FR |
| *35S:MYB68-mCherry* | *p35S* | B3 | MYB68 | m-Cherry | E1 | FR |
| *35S:MYB36-mCherry* | *p35S* | B3 | MYB68 | m-Cherry | E1 | FR |

**Table S5 Oligos used in this study**

| **Element** | **Primer for cloning** | **Sequence** |
|---|---|---|
| **pMYB68** | A-pMYB68 F | AACAGGTCTCAACCTAATACGATGCTACTCTGTTGTT |
| | Br-pMYB68 R | AACAGGTCTCATGTTAGGAGGATGGTGTATGATAATG |
| **pMYB84** | A-pMYB84 F | AACAGGTCTCAACCTCGTGGACTTGGACTTGTTTA |
| | Br-pMYB84 R | AACAGGTCTCATGTTACTTGTACTCCTAGTGAAGTCTTG |
| **pMYB53** | A-pMYB53 F | AACAGGTCTCAACCTTGACCTCCTGGATTTCAAGAA |
| | Br-pMYB53 R | AACAGGTCTCATGTTGTTTGATCAACTTCTGTATCTAACAA |
| **pMYB37** | Br-pMYB84 R | AACAGGTCTCAACCTGTACTTCACATGATAACACG |
| | Br-pMYB37 R | AACAGGTCTCATGTTTCTCGTTAGTGAATTGAAG |
| **pGPAT5** | A-pGPAT5 F | AACAGGTCTCAACCTTCGCAAACGTCAATGGTCTAT |
| | Br-pGPAT5 R | AACAGGTCTCATGTTTTCTTTTGTTTTTTGCTCGAATATTA |
| **MYB68** | C-MYB68 F | AACAGGTCTCAGGCTCAATGGGAAGAGCACCGTGTTGT |
| | Dr-MYB68 R | AACAGGTCTCACTGACACATGATTTGGCGCATTGAA |
| **MYB36** | C-MYB36 F | AACAGGTCTCAGGCTCAATGGGAAGAGCTCCATGCT |
| | Dr-MYB36 R | AACAGGTCTCACTGAAACACTGTGGTAGCTCATCTGAG |

| | | |
|---|---|---|
| **GELP51** | C-GELP51 F | AACAGGTCTCAGGCTCAATGGCCACAAGAGCTTCTA |
| | Dr-GELP51 R | AACAGGTCTCACTGATCACATATCTCTAAGTTTGC |
| **pMYB38** | pMYB38 F | GAAAAGTTGGGTACCGATAGGGGAGGCACCTATTTGAG |
| | pMYB38 R | ACAAACTTGCCCGGGCTCTCTATCTGTCTCTCTTGG |
| **pMYB87** | 1/2B1pMYB87 F | AAAAAGCAGGCTGGGTATAAGTACAGCCCAAGT |
| | 1/2B2pMYB87 R | AGAAAGCTGGGTATCTTGTTCTCTAGGTATTTATC |
| **gRNA MYB87** | MYB87gRNA F | ATTGGAAAGGGCCATGGTCGACGG |
| | MYB87gRNA R | AAACCCGTCGACCATGGCCCTTTC |
| **gRNA MYB68** | MYB68gRNA F | AATAGTGGCACAGGGTTTTAGAGCTAGAA |
| | MYB68gRNA R | TGCCACTATTCTCGATCAATCACTACTTCGAC |
| **pGPAT5** | A-pGPAT5F | AACAGGTCTCAACCTTCGCAAACGTCAATGGTCTAT |
| | Br-pGPAT5 R | AACAGGTCTCATGTTTTCTTTTGTTTTTTGCTCGAATATTA |
| **pGELP8** | A-pGELP8F | AACAGGTCTCAACCTACCATATGCGAATCCTCCTT |
| | Br-pGELP8 R | AACAGGTCTCATGTTGGGAGAGGCCGGAAGG |
| **pGELP51** | A-pGELP51F | AACAGGTCTCAACCTGATGATGATGAAGATGATAC |
| | Br-pGELP51 R | AACAGGTCTCATGTTATTTCTTGATTTTGAGAAATG |
| **pGELP38** | A-pGELP38F | AACAGGTCTCAACCTTGTTGTTTGAGGAGGAGTTC |
| | Br-pGELP38 R | AACAGGTCTCATGTTGTCGCCAAATTTGAATGTGTG |
| **pCASP2** | A-pCASP2F | AACAGGTCTCAACCTATAACCAGCAAATCGAG |
| | A-pCASP2 BSAI | AACAGGTCTCATAGGTGGGACCCAATGAAAGA |
| | Br-pCASP2 BSAI | AACAGGTCTCACCTATGGGATTAAGCTTA |
| | Br-pCASP2 R | AACAGGTCTCATGTTTTTCTTTACTTTGTTT |
| **NnLUX** | C-NnLUX | ATACAGGTCTCAGGCTATATGCGTATAAATATCTCTCTGTCTAGC |
| | Dr-NnLUXR | ATAAAGGTCTCGCTGATCATTTGGCATTCTCGACGATTTTAC |

| Allele | Primer for genotyping | Sequence |
|---|---|---|
| **myb84-1** | WT : LP+ RP<br>Mut: RP+ LB | LP ACTTTTTGACATCTCCCCCTC<br>RP GTGTTGGCGCTCAAGATAGAG<br>LB ATTTTGCCGATTTCGGAAC |
| **rax1-3** | WT : LP + RP<br>Mut: RP + LB | LP TCGGACATTTCAGTTTGGAAG<br>RP TGTGAAAAGACCAACCTCACC<br>LB ATTTTGCCGATTTCGGAAC |
| **rax2-1** | WT : LP + RP<br>Mut: LP + JL | LP AAAAAGCAGGCTTTATGGGTAGGGCTCCATGTT<br>RP AGAAAGCTGGGTTTCAGTAGTACAACATGAACTTATC<br>JL CATTTTATAATAACGCTGCGGACATCTAC |
| **rax3-1** | WT : LP + RP<br>Mut: LP + En | LP ACTTTTTGACATCTCCCCCTC<br>RP GTGTTGGCGCTCAAGATAGAG<br>En AGAAGCACGACGGCTGTGATAGGA |
| **wox4-1** | WT : LP + RP<br>Mut: GK + RP | LP GCTCGTAGGTCAATGTCAATCTC<br>RP CCTATCTGTTCTTGAGTCGGG<br>GK ATATTGACCATCATACTCATTGC |
| **bp-9** | WT : BP14 + BP3<br>Mut: BP14 + DSMP1 | BP14: TGTTAAGGGTTAGAACACCATG<br>BP3: GACAACAGCACCACTCCTCAAA<br>Dspm1: CTTATTTCAGTAAGAGTGTGGGGTTTTGG |
| **myb68-2-4** | PCR: F+R<br>Seq: S | F:AACAGGTCTCAGGCTCAATGGCCACAAGAGCTTCTA<br>R: AACAGGTCTCACTGATCACATATCTCTAAGTTTGC<br>S: TGCAACTCTTCCCACATCTCC |
| **myb87-3** | PCR: F+R<br>Seq: | F: GGTTAGAGGTGCATGGTGGA<br>R :AAATAGCTCGGCCTGGAAAT<br>S: TCATTGCCTTTTTCTCACCA |

| GENE | Reference | Sequence |
|---|---|---|
| FAR4 | Based on (Andersen et al. 2021) | F CATGCAACGGTTTCACAGTGAGGT<br>R TGAGCGGTCCAAATGTGTTGACG |
| HORST | Based on (Andersen et al. 2021) | F GGAGACACGTGGCAATGATCAGGA<br>R GTCGAGCCTCTTGGCACGAAAGT |
| GPAT5 | Based on (Andersen et al. 2021) | F ACCGCCGTGTTAATTTGTTTGTTGG<br>R CCGTCGTGAAATATCACCGGAAGT |
| GELP8 | This study | F TGGTCACTTCTGTGAACTCG<br>R TTAGGATCCGAGAGACCAAG |
| GELP38 | Based on (Ursache et al. 2021) | F GTTTGGTGAAGCCTATGACCTCGTC<br>R CATCGGCACGCATGAAGTCAAATC |
| GELP51 | Based on (Ursache et al. 2021) | F TGTCCATGCAAACGTCTACGATCTC<br>R AAGTCGGTCCGCACGGAATTATAC |
| GELP96 | Based on (Ursache et al. 2021) | F CCGGAGCTAAATTCTCTTTCGCAGA<br>R TAGCCGAATCGGACGGATGAAATG |
| MYB41 | This study | F GTACGCTCCCCAAAAATGCTGGAC<br>R GCGGCTATTGCTGACCACTTGTTTC |
| MYB39 | Based on (Shukla et al. 2021) | F CCCTCCTTGGCAACAAATGGTC<br>R GATCGTTGGTTCTTGGCTCGTG |
| MYB53 | Based on (Shukla et al. 2021) | F TGCGGTTCTAGGCAACAAGTGG<br>R TCATCGGTTCTTGGCTGATGGG |
